# Stable Maintenance of Two-Cell-Like Cells from Embryonic Stem Cells Reveals Chromatin and Super Enhancer Regulation of MERVL Elements

**DOI:** 10.1101/2025.09.25.678376

**Authors:** Rui Geng, Benjamin L. Kidder

**Affiliations:** Department of Oncology, Wayne State University School of Medicine, Detroit, MI, USA; Karmanos Cancer Institute, Wayne State University School of Medicine, Detroit, MI, USA

**Keywords:** embryonic stem cell, two-cell, 2CLC, pluripotency, totipotency, MERVL, LTR, chromatin, epigenetics, super enhancer, RNA-Seq, ChIP-Seq

## Abstract

Mouse embryonic stem cells (ESCs) occasionally transit into a rare two-cell-like (2C) state characterized by transient activation of endogenous retroviruses such as MERVL and expression of 2C-specific genes including the Zscan4 cluster. These 2C-like cells (2CLCs) resemble early blastomeres and display expanded developmental potential, but their unstable and sporadic nature has hindered mechanistic studies. Here, we establish stable 2CLCs (s2CLCs) that maintain persistent MERVL expression and homogeneous 2C gene activation. Live-cell imaging revealed uniform and sustained MERVL activity in s2CLCs, contrasting with the heterogeneous and transient expression observed in conventional ESCs. Transcriptome profiling demonstrated robust induction of 2C-specific regulatory networks, and embryoid body differentiation combined with machine learning uncovered increased lineage variability and expanded developmental trajectories. Epigenomic profiling further revealed unique chromatin states, distinctive super enhancer landscapes, and active enhancer marking at MERVL loci. Together, these findings demonstrate that stable maintenance of the 2C-like state is achievable in vitro, providing a powerful model to dissect ERV-driven transcriptional regulation, epigenomic remodeling, and totipotent-like developmental potential.

## INTRODUCTION

Mouse embryonic stem cells (ESCs), derived from the inner cell mass (ICM) of blastocyst-stage embryos, are pluripotent and capable of generating derivatives of all three embryonic germ layers. In contrast, totipotent cells such as zygotes and two-cell (2C) stage blastomeres can give rise to both embryonic and extraembryonic lineages^1^. Understanding how cells transition into the totipotent state is a central question in developmental biology, as it defines the earliest developmental potential. In mice, zygotic genome activation (ZGA) peaks at the 2C stage^2^, coinciding with robust expression of repetitive sequences such as major satellites^3, 4^, endogenous retroviral elements (ERVs) including MERVL^4–6^, and numerous 2C-specific genes^7–9^.

ESC cultures occasionally generate rare two-cell-like cells (2CLCs) that transiently resemble blastomeres and co-express MERVL with genes such as Zscan4^6, 10, 11^. These cells revert rapidly, and MERVL expression is silenced beyond the 2C stage^12, 13^. Many 2C-specific genes are directly driven by MERVL long terminal repeat (LTR) promoters, highlighting a functional role for retroelements in totipotent-like gene activation. Transposable elements account for more than half of the mammalian genome, with ERV-derived LTR retrotransposons representing a major class^14^. While ESCs are characterized by transcriptional and epigenetic heterogeneity^15–17^, evidence suggests that epigenetic mechanisms such as histone modifications contribute to MERVL activation. MERVL-positive ESCs display elevated histone acetylation^4, 18^, and perturbations that alter chromatin states can increase or decrease the frequency of MERVL-expressing cells^4, 18–26^. These findings link chromatin remodeling to ERV activity, yet mechanistic analysis has been hindered by the fleeting nature of the 2C-like state.

A major barrier in the field has been the inability to stably capture a homogeneous population of 2CLCs without extensive perturbation of the epigenome. As a result, the transcriptional and chromatin mechanisms sustaining totipotent-like identity have remained incompletely defined. We hypothesized that establishing stable 2CLCs (s2CLCs) would overcome this limitation by enabling long-term analysis of MERVL-driven transcription and chromatin states. Here, we report the successful maintenance of ESCs in a stable 2C-like state, characterized by persistent expression of MERVL and 2C-specific genes. Transcriptome and epigenome profiling revealed a distinct regulatory network, expanded differentiation potential, and unique enhancer landscapes marked by activating histone modifications and super enhancer formation. The establishment of s2CLCs provides a tractable model for dissecting ERV-driven transcriptional regulation, epigenomic remodeling, and totipotent-like developmental potential.

## RESULTS

### Identification and prospective isolation of stable 2C-like cells by MERVL activity

To visualize and track the 2C-like state in ESCs, we generated stable integrations of a MERVL promoter–driven 2C::tdTomato reporter^4^ (**Fig. 1A**; See Methods). Flow cytometry of bulk cultures revealed ∼15–16% reporter-positive cells (**Fig. S1**), consistent with the presence of a rare 2C-like subpopulation under self-renewal conditions. We then used FACS to prospectively isolate 2C::tdTomato+ cells and clonally expand reporter-positive colonies.

**Figure 1.**
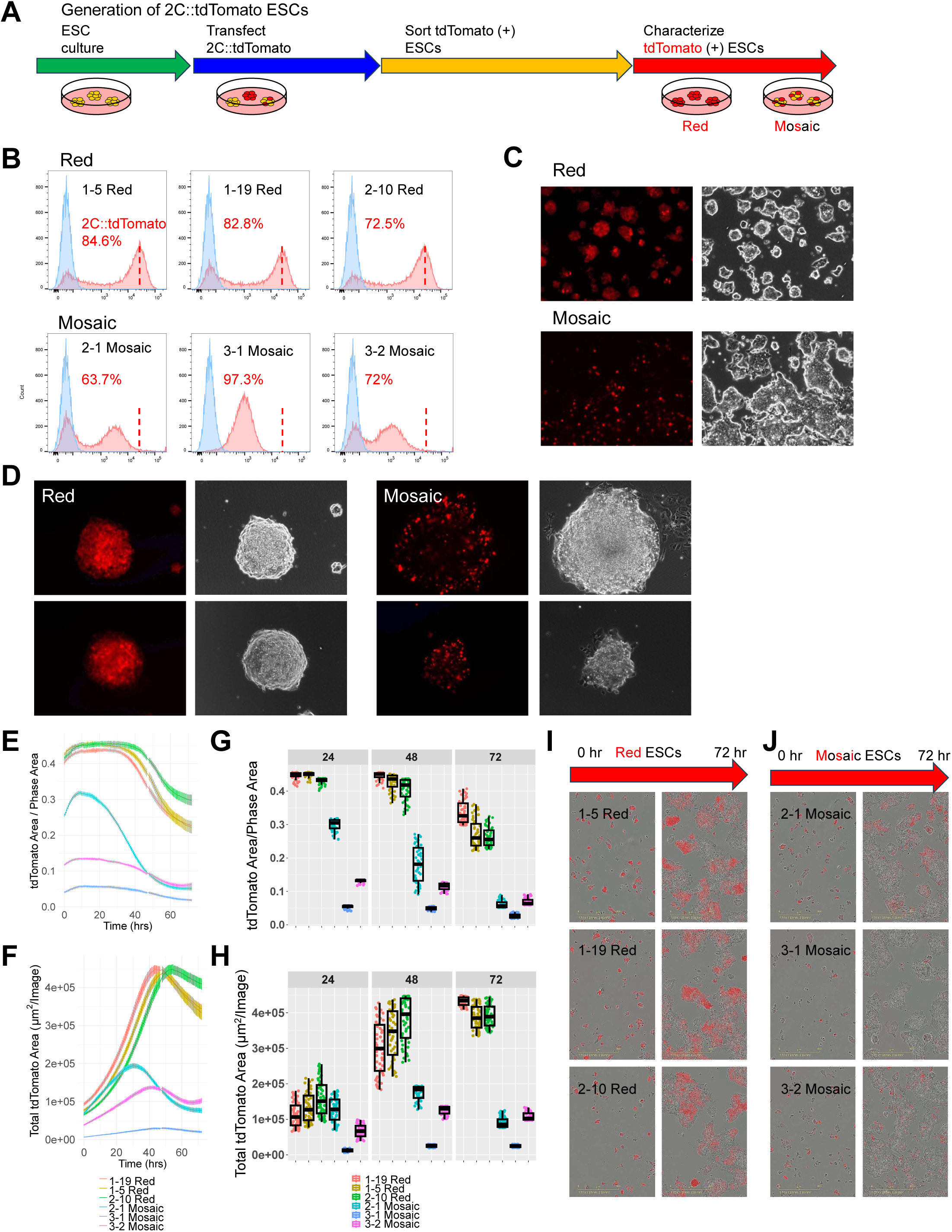
Identification and characterization of stable MERVL-positive 2C-like cells. (**A**) Schematic of the experimental workflow. ESCs were transfected with a modified 2C::tdTomato reporter vector and subjected to FACS sorting to isolate MERVL-positive cells. The resulting populations carried stable MERVL promoter–driven tdTomato integrations, enabling visualization and prospective isolation of 2C-like cells. (**B**) The 2C::tdTomato reporter was used to detect MERVL expression, enabling fluorescence-activated cell sorting (FACS) of 2C-like cells (2CLCs) from ESCs. Red (top) and mosaic (bottom) ESC colonies contained a higher proportion of MERVL-positive cells compared with conventional ESCs (see **Fig. S1**). (**C**) Post-sort analysis of ESCs showed variable MERVL-positivity in red and mosaic populations. Fluorescent (2C::tdTomato; left) and bright field (right) images are shown. (**D**) Representative fluorescent and bright field images of red (left) and mosaic (right) MERVL-positive ESCs. (**E–H**) Sequential imaging of red and mosaic stable 2C-like cells (s2CLCs) under self-renewal conditions, with quantification of tdTomato area relative to phase area or total tdTomato area (µ²/image). These analyses show that red s2CLCs maintain a higher and more stable fraction of MERVL-positive cells compared with mosaic s2CLCs during the initial 48 hours. After 48 hours, 60–70% of cells were reporter-positive in fluorescent images. (**I, J**) Representative bright field and fluorescent images at 0 and 72 hours, demonstrating sustained MERVL expression in red s2CLCs compared with the more sporadic expression observed in mosaic s2CLC colonies.

Two reproducible phenotypic classes emerged among reporter-positive colonies: (i) “mosaic” colonies with heterogeneous, generally lower reporter signal, and (ii) “red” colonies with uniformly higher reporter intensity. Flow cytometry of expanded clones confirmed that both classes maintained substantial reporter-positive fractions, with red clones exhibiting markedly higher MERVL reporter levels (**Fig. 1B–C**). In some red clones, up to 84% of cells were reporter-positive, far exceeding conventional ESCs, whereas mosaic clones contained fewer reporter-positive cells and at lower intensity (**Fig. 1B–C**). High-magnification micrographs illustrate the contrasting expression patterns (**Fig. 1D**).

Live-cell imaging over 72 h showed that MERVL activation in mosaic colonies was dynamic, with transitions into and out of the positive state, whereas red colonies displayed sustained, homogeneous reporter expression. Quantification by total tdTomato area per image and tdTomato/phase area ratio demonstrated that red s2CLCs retained higher and more stable reporter levels throughout the time course (**Fig. 1E–H**). After ∼40–48 h, 60–70% of s2CLCs were reporter-positive, and by 72 hrs. most cells in red clones expressed the reporter, while mosaic clones remained lower on average (**Fig. 1E–H**). Representative fields at 0 h and 72 h are shown for both classes (**Fig. 1I–J**). Based on their persistence and uniformity, we define these red and mosaic reporter-positive lines collectively as stable 2C-like cells (s2CLCs).

### Transcriptome profiling reveals a distinct 2C program in red s2CLCs

To characterize the expression profiles of red and mosaic MERVL-positive ESCs, and to understand the expression of 2C-genes, ERVs, and differentially expressed genes between red and mosaic s2CLCs and conventional ESCs, we performed transcriptome analysis using RNA-Seq. Conventional ESCs (+LIF) and differentiated ESCs (–LIF) were used as controls (see Methods). edgeR differential expression analysis revealed variations in the number of upregulated and downregulated genes in red and mosaic ESCs (**Fig. 2A**). Principal component analysis (PCA) demonstrated that red s2CLCs are more distinct from conventional ESCs along the PC1 axis, which accounts for most of the variability, compared to mosaic s2CLCs (**Fig. 2B**).

**Figure 2.**
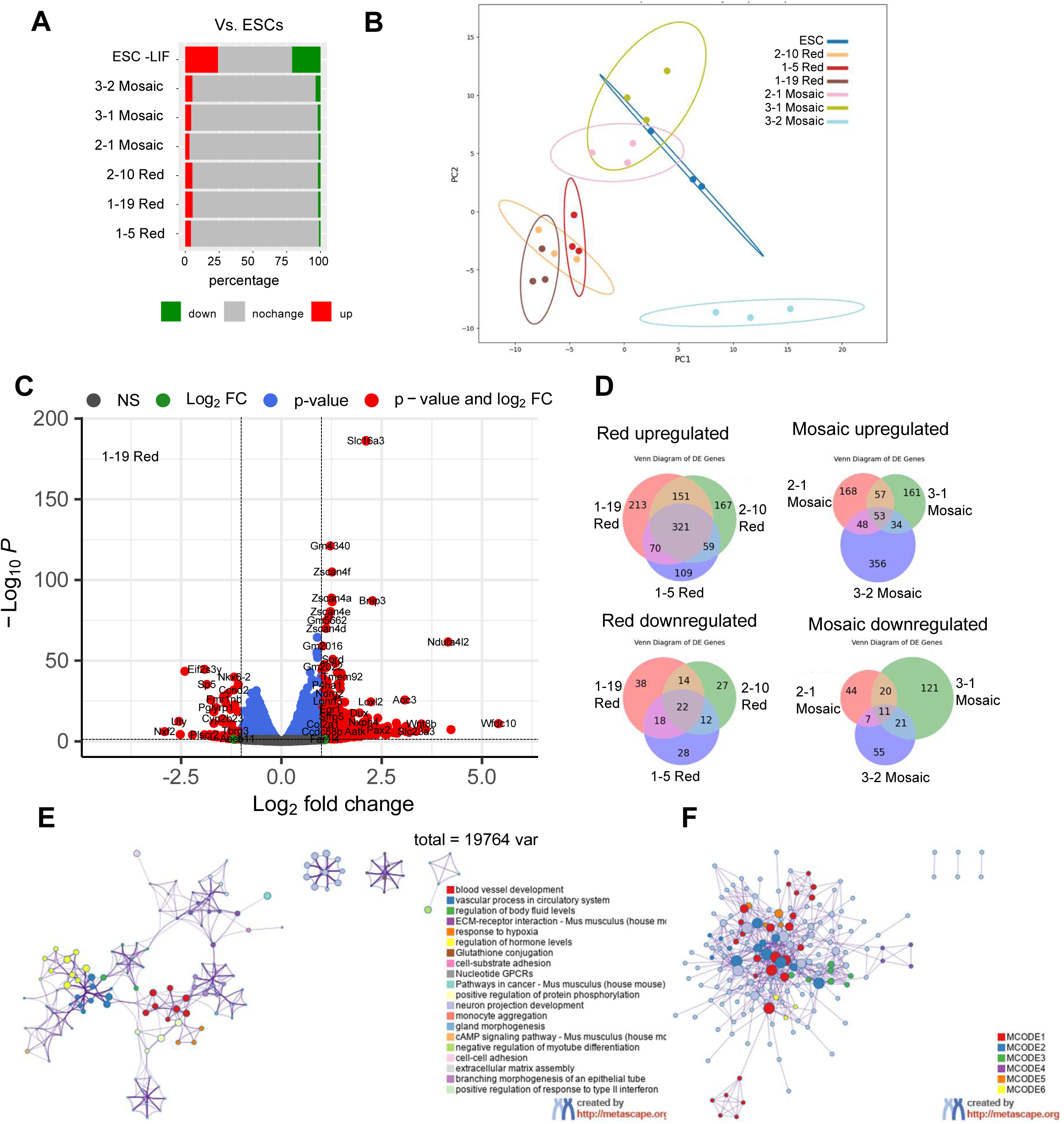
Transcriptome analysis of s2CLCs. (**A**) Differential expression analysis of MERVL-positive ESCs and differentiated ESCs compared with conventional ESCs, showing the number of upregulated and downregulated genes identified by edgeR. (**B**) Principal component analysis (PCA) illustrating that red s2CLCs cluster distinctly from conventional ESCs, whereas mosaic s2CLCs display greater variability and remain closer to conventional ESCs. (**C**) Volcano plot of differentially expressed genes between a representative red s2CLC line (1-19 red) and conventional ESCs. Each point represents a gene, plotted by log2 fold change (x-axis) and significance (–log10 p-value, y-axis). Grey points denote non-significant genes, green points represent significant fold changes, blue points indicate significant p-values, and red points highlight genes significant by both criteria. Elevated expression of transcription factors and other 2C-associated regulators is observed in red s2CLCs. (**D**) Genes upregulated (upper panels) and downregulated (lower panels) in red s2CLCs (left) and mosaic s2CLCs (right). Venn diagrams show unique and shared upregulated genes across red and mosaic populations. (**E**) Cytoscape visualization of Metascape enrichment analysis, depicting gene ontology clusters specifically activated in red s2CLCs. (**F**) Protein interaction networks identified by Metascape, illustrating biochemical assemblies and signaling pathways enriched in red s2CLCs. Notable transcriptional regulators include Stat3, among others. Nodes are color-coded by gene ontology enrichment p-value, and protein-protein interaction clusters are annotated in corresponding colors.

Volcano plots were used to visually identify and track genes that were upregulated and downregulated in a pairwise comparison. Red s2CLCs exhibited elevated expression of genes such as Zscan4d^7^, Dux^8, 9^, Zim3, Wnt3, Wnt8b, and Trim56 (**Fig. 2C; Fig. S2**). Notably, Zscan4a and Dux are two key transcription factors that regulate the 2C-like state^7–9^.

We identified 321 genes that were upregulated in red s2CLCs compared to conventional ESCs. In contrast, mosaic s2CLCs exhibited an upregulation of only 53 genes relative to conventional ESCs. Additionally, there were only 22 and 11 genes downregulated in red and mosaic ESCs, respectively (**Fig. 2D**).

MA plots further highlighted statistical characteristics of differentially expressed (DE) genes (fold-change and p-value; **Fig. S3**). Gene ontology (GO) functional annotation of DE genes using clusterProfiler^28^ allowed for the comparison of GO terms enriched in activated or suppressed genes in red and mosaic ESCs (**Fig. S4**). Specifically, pathways that were activated include transmembrane signaling receptor activity, positive regulation of cell adhesion, regulation of cell–cell adhesion, calcium ion binding, and signaling receptor activity, which were prominently activated in red s2CLCs. This suggests that red s2CLCs have heightened activities in cell adhesion, ion transport, and cell signaling.

Additional functional genomics analysis of regulated genes in red s2CLCs was conducted through Metascape^29^ enrichment analysis. Enrichment networks were displayed using Cytoscape^30^ to highlight GO clusters (**Fig. 2E**). Additionally, protein network analysis was performed using Metascape, involving the assessment of gene lists within the framework of protein interactions to identify biochemical assemblies or signaling pathways that influence biological outcomes (**Fig. 2F**). The Metascape-MCODE algorithm was further used to independently identify protein complexes within the expansive network. This analysis uncovered a wide range of transcription factors expressed at elevated levels in red ESCs. TRRUST analysis using Metascape also shed light on targets of TFs whose expression is enriched in red ESCs (**Fig. S5**). Stat3, a known pluripotency regulator in ESCs, had multiple targets identified. Additionally, gene expression of targets for Ets1, Sp1, Etv2, Fos, Trp53, and Pparg were found to be enriched in red ESCs compared to conventional ESCs.

### Expression of endogenous retroelements in stable 2CLCs

To further profile the expression of repetitive DNA elements in red and mosaic s2CLCs, we used RepeatMasker annotations of repeat family members and repeat names, and performed edgeR analysis to identify differentially expressed (DE) elements relative to conventional ESCs. We observed several ERVL-class retrotransposons consistently upregulated in red s2CLCs, including MERVL-int, MT2_Mm, RLTR6_Mm, ORR1A3-int, and GSAT_MM, whereas mosaic s2CLCs showed significant upregulation of only RLTR13B2. These findings, illustrated in volcano plots **(Fig. S6A)** and heatmaps **(Fig. S6B–C)** for both repeat families and ERVL elements, confirm that red s2CLCs sustain elevated expression of MERVL-int and MT2_Mm. Importantly, the strong induction of RLTR6_Mm, ORR1A3-int, and GSAT_MM in red s2CLCs represents an expanded set of ERV-associated elements not previously linked to the 2C-like state. Conversely, several repetitive elements, including MER68-int, MLT1F1-int, Tigger9a, and U4, were consistently downregulated in red s2CLCs.

### Expanded differentiation of s2CLCs revealed through embryoid body assays

2CLCs have an expanded differentiation potential, enabling contributions to both embryonic and extraembryonic lineages. To evaluate whether this feature is preserved in stable 2CLCs, we performed embryoid body (EB) differentiation assays over a 12-day period, which mimic early embryonic development. Morphological analysis of the resulting EBs revealed that s2CLCs displayed greater heterogeneity compared to conventional ESCs, as demonstrated by increased variability in EB shapes (**Fig. 3A**).

**Figure 3.**
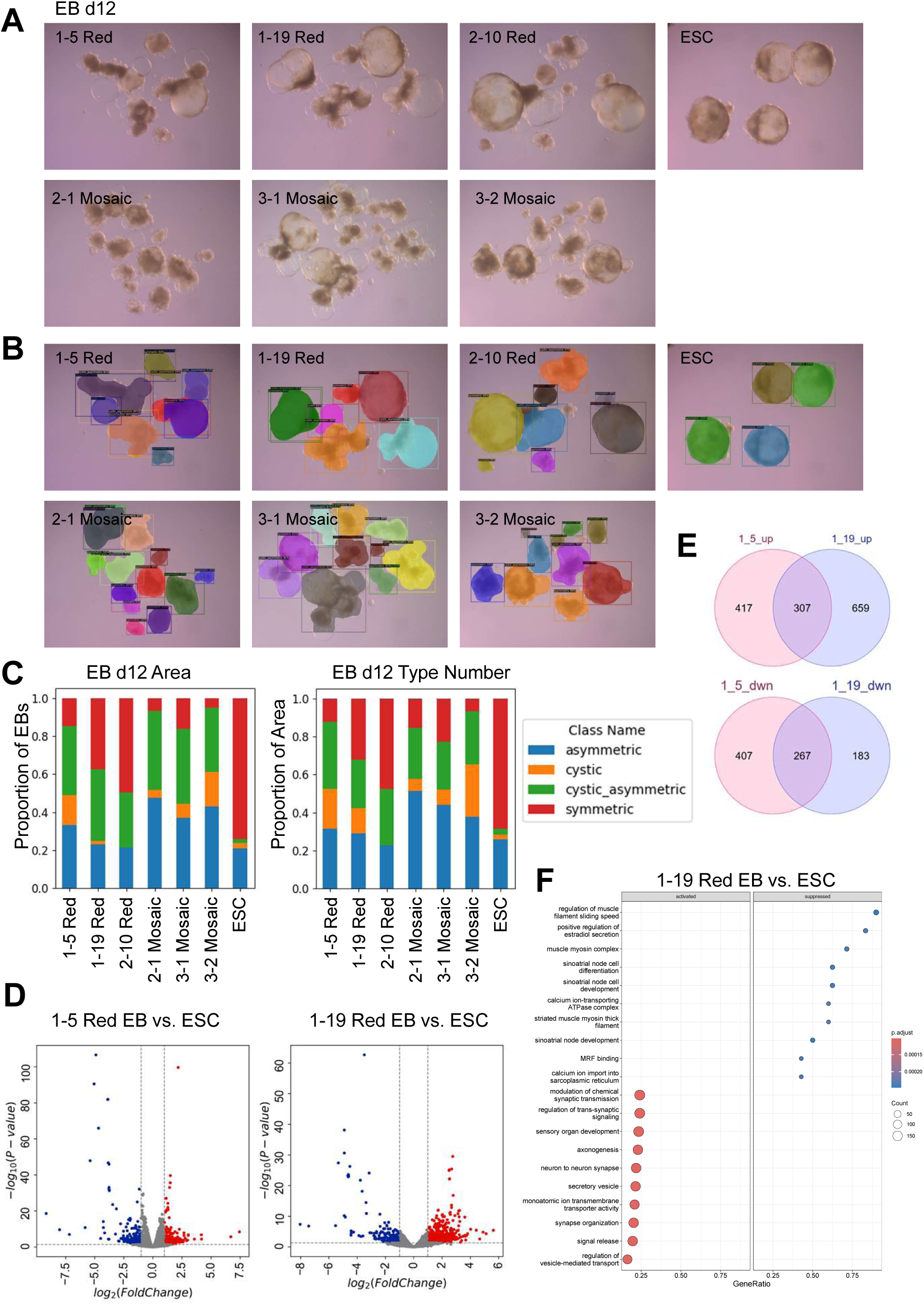
Differentiation and transcriptome analysis of s2CLCs. **(A)** Bright field microscopy of red and mosaic s2CLCs, and conventional ESCs, differentiated into embryoid bodies (EBs) without LIF for 12 days. (**B**) Machine learning– based visualization of predicted EB morphologies, categorized as symmetric, asymmetric, cavitated, and cavitated-asymmetric. (**C**) Image segmentation results shown as stacked bar plots, illustrating the distribution of EB morphologies by area and number. (**D**) Volcano plot of differentially expressed genes comparing day 12 EB-differentiated red s2CLCs (clones 1-5 and 1-19) with conventional EBs. Genes are plotted by log2 fold change (x-axis) and significance (–log10 p-value, y-axis). Grey points represent non-significant genes, green points indicate significant fold changes, blue points denote significant p-values, and red points highlight genes significant by both criteria. (**E**) Venn diagram showing the overlap of upregulated and downregulated genes between red s2CLCs (clones 1-5 and 1-19) and conventional EB-differentiated ESCs. (**F**) Gene Ontology enrichment analysis (clusterProfiler) of upregulated and downregulated genes in day 12 EB-differentiated red s2CLCs (clone 1-19) compared with conventional EB-differentiated ESCs.

To quantify these variations, we applied a machine-learning–based image segmentation approach to classify EB morphologies into four categories: symmetric, asymmetric, cavitated, and cavitated-asymmetric (Methods). Segmentation highlighted distinctive features of red and mosaic s2CLC-derived EBs compared to controls (**Fig. 3B; Fig. S7A– B**). Notably, both red and mosaic s2CLCs produced fewer and smaller symmetric EBs but a greater proportion of cystic and cystic-asymmetric EBs (**Fig. 3C**). The predominance of cystic structures is consistent with trophectodermal lineage potential, aligning with the expanded potency of 2C-like states.

To probe differentiation trajectories at the transcriptional level, we performed RNA-seq after 12 days of EB differentiation in the absence of LIF. Red s2CLC-derived EBs exhibited 307 upregulated and 267 downregulated genes relative to conventional ESC-derived EBs (**Fig. 3D–E**). GO enrichment analysis (clusterProfiler) identified biological processes enriched among upregulated genes, including pathways related to muscle and cardiac development, calcium handling, and contractile fiber organization (**Fig. 3F; Fig. S8**). Together, these findings demonstrate that s2CLCs undergo altered and broadened differentiation programs, consistent with their expanded lineage potential.

### Epigenome profiling of red and mosaic s2CLCs

We next examined whether stable 2CLCs are accompanied by distinctive chromatin landscapes. To do so, we analyzed H3K4me3^31^ (active promoters/TSS^32–34^) and H3K27ac^35^ (active enhancers/super enhancers) by ChIP-seq.

We first quantified global peak numbers across red and mosaic s2CLCs, conventional ESCs, and differentiated ESCs. H3K4me3 peak counts were heterogeneous but consistently elevated in red and mosaic s2CLCs compared to ESCs and differentiated cells, ranging from ∼29,857–37,727 in red clones (**Fig. 4A, top**). H3K27ac peaks also increased, ranging from ∼33,663–39,794 in red s2CLCs (**Fig. 4A, bottom**). Mosaic clones tended to show slightly higher counts, consistent with a more variegated transcriptional landscape. In contrast, differentiated ESCs showed reduced H3K4me3, highlighting the loss of promoter activity upon exit from pluripotency.

**Figure 4.**
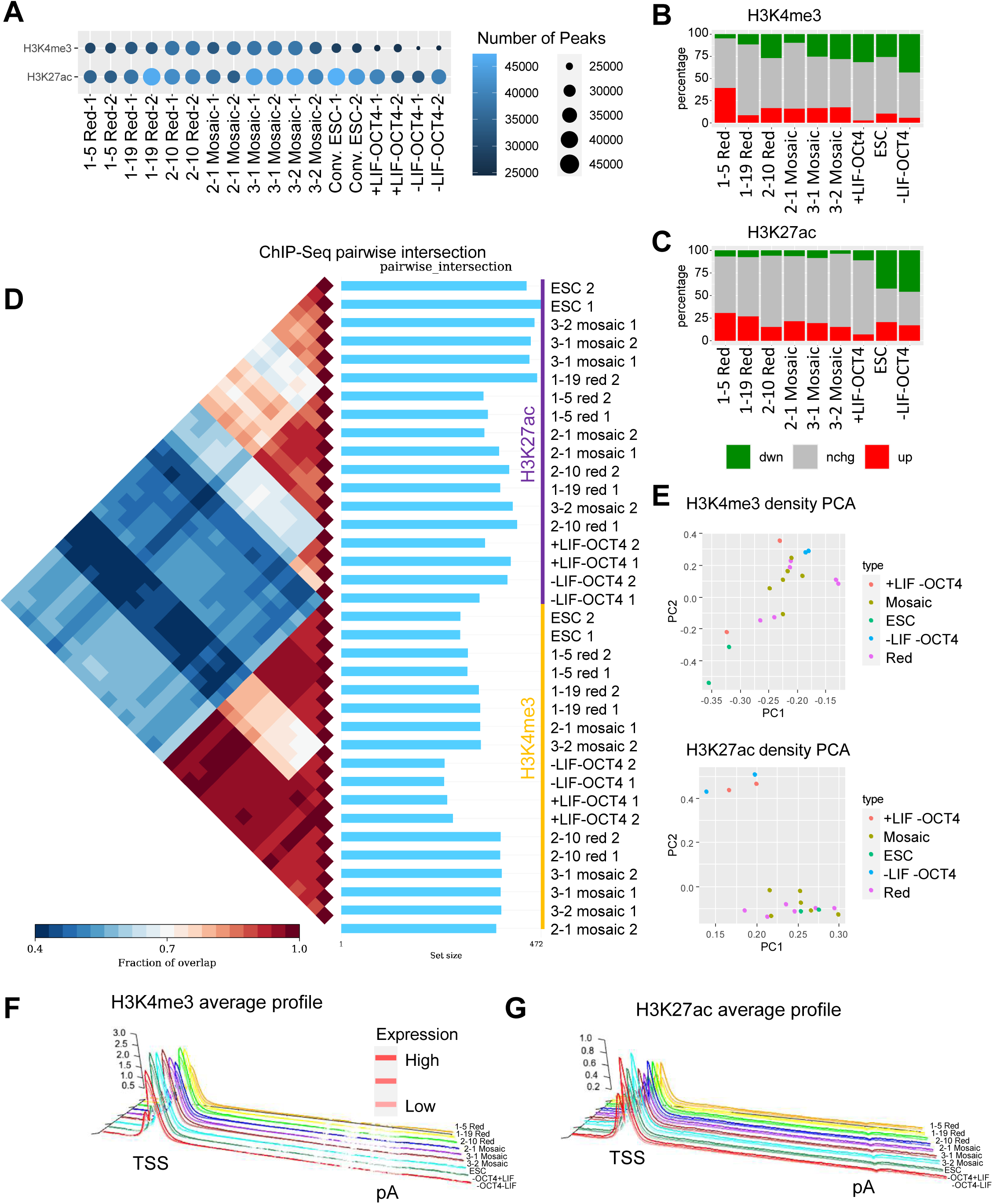
Comparative epigenetic profiling of histone modifications in s2CLCs across differentiation states. (**A**) Bubble plot showing the number of ChIP-enriched peaks for H3K4me3 and H3K27ac, with bubble size and color indicating peak counts. (**B, C**) Percentage change in (**B**) H3K4me3 and (**C**) H3K27ac levels across experimental conditions. Red indicates increased peaks and green indicates decreased peaks, as determined by SICER-compare analysis of multiple samples. These comparisons reveal differential epigenetic regulation in red and mosaic s2CLCs relative to conventional ESCs. (**D**) Pairwise comparison of ChIP-enriched peak overlap performed using Intervene. (**E**) Principal component analysis (PCA) of normalized H3K4me3 and H3K27ac tag densities. (**F, G**) Average density profiles of (**F**) H3K4me3 and (**G**) H3K27ac across all RefSeq genes, spanning transcription start sites (TSS) to polyadenylation sites (pA). Genes were stratified into quartiles based on expression levels in conventional ESCs, illustrating global patterns of histone modifications and the distinctive epigenetic landscapes of red and mosaic s2CLCs.

To systematically compare signal dynamics, we applied SICER-compare (FDR < 0.001, FC > 1.5)^36^. Red and mosaic s2CLCs showed significant increases in H3K4me3 and H3K27ac peaks relative to ESCs, whereas –LIF/–OCT4 differentiation caused pronounced losses (**Fig. 4B**). For example, “1-5 red” clones exhibited increased H3K4me3, while – LIF/-OCT4 cells had decreased H3K4me3. Two red clones also showed elevated H3K27ac compared to ESCs (**Fig. 4C**), suggesting enhancer hyperactivation.

Peak intersections using Intervene^37^ showed that red and mosaic s2CLCs shared large fractions of H3K4me3/H3K27ac peaks, clustering more closely with one another than with ESCs or differentiated states (**Fig. 4D**). PCA of H3K27ac densities across merged peaks clearly separated s2CLCs from ESCs and differentiated cells (**Fig. 4E**), indicating that enhancer acetylation better discriminates the s2CLC state than H3K4me3.

Metagene profiles of H3K4me3 and H3K27ac across TSS to polyA regions further illustrated globally elevated activation marks in s2CLCs (**Fig. 4F–G**). HOMER^38^ annotation confirmed enrichment of both marks in promoters, introns, and intergenic regions (**Fig. S9A–B**). deepTools^39^ k-means clustering identified subsets of genes with broad H3K4me3 domains and distinct H3K27ac enrichment at highly expressed loci (**Fig. S9C–D**). Density comparisons against ESCs confirmed broad promoter and enhancer activation in red and mosaic s2CLCs (**Fig. S9E–F**).

### Enhancer remodeling in s2CLCs: ERV-proximal activity and super enhancers

Given the central role of ERVs in 2C biology, we next asked whether enhancer activation was linked to MERVL loci. deepTools density analysis revealed that both MERVL-int and MT2_Mm were strongly enriched for H3K27ac in red s2CLCs, but only weakly marked in mosaic s2CLCs, ESCs, or differentiated cells (**Fig. 5A–B**). This indicates that stable ERV activation in red s2CLCs is reinforced by proximal enhancer acetylation, potentially stabilizing MERVL-driven transcription.

**Figure 5.**
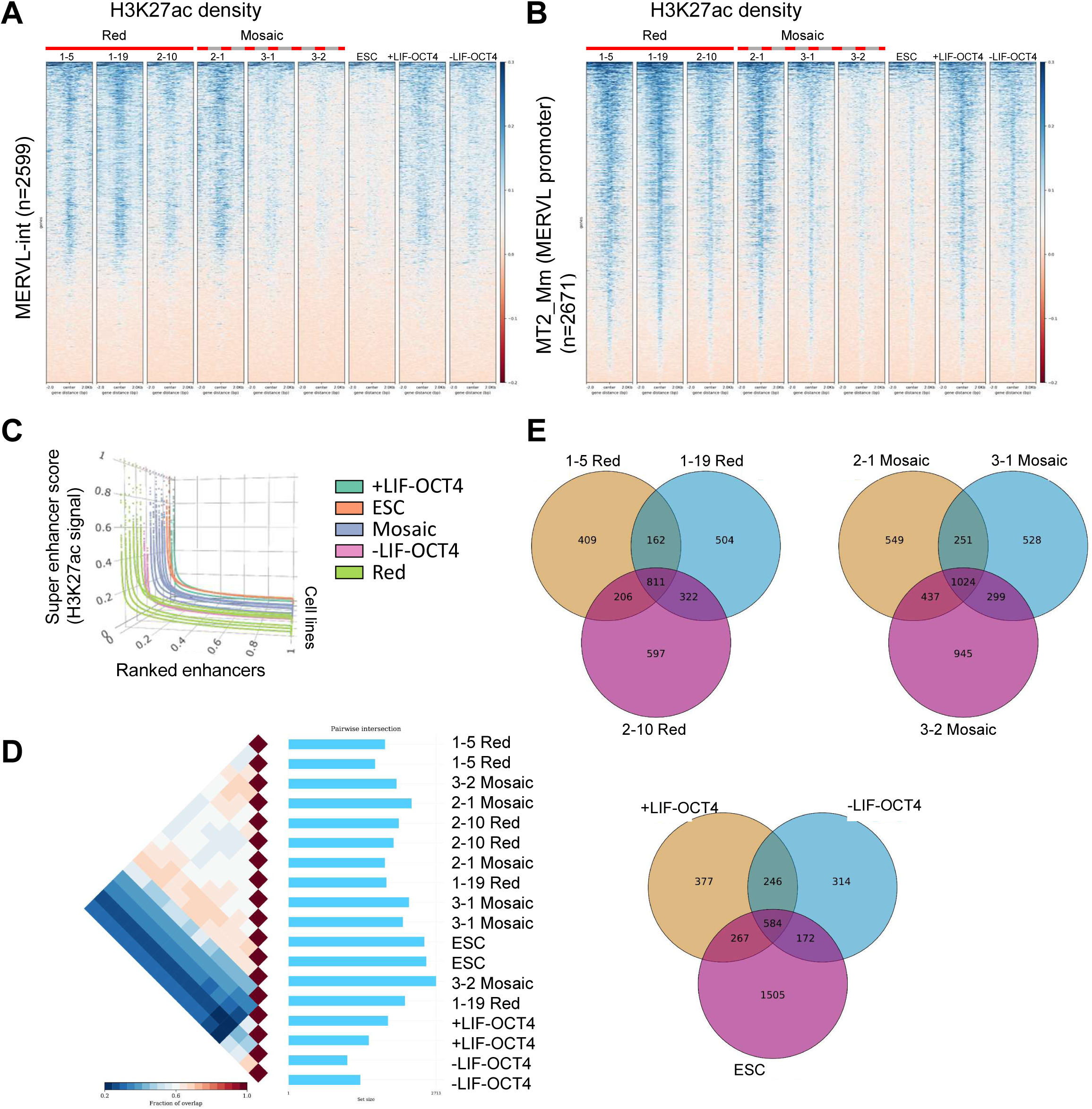
Differential enrichment of H3K27ac and super enhancer analysis of s2CLCs. (**A, B**) Distribution of H3K27ac signals around (**A**) MERVL-int and (**B**) MT2_Mm (MERVL promoter) repeat elements, generated using deepTools. (**C**) Saturation curve of H3K27ac densities in s2CLCs and ESCs, showing the distribution of typical enhancers and super enhancers (SEs) ranked by H3K27ac density. Normalized H3K27ac ChIP-seq signals are plotted across enhancer regions. Super enhancers were identified using HOMER (see Methods) and defined as regions with a slope >1. (**D**) Pairwise comparison of super enhancer overlap across s2CLCs and ESCs under different culture conditions, performed with Intervene. (**E**) Venn diagrams illustrating unique and shared super enhancer regions in red and mosaic s2CLCs.

To broaden this view, we identified super enhancers (SEs) by ranking stitched H3K27ac peaks (HOMER^38^ –superSlope –1000; merge window 12.5 kb). Red s2CLCs displayed high-intensity SEs distinct from mosaic s2CLCs, ESCs, and differentiated cells (**Fig. 5C**). Overlap analysis using Intervene revealed limited sharing between groups, underscoring cell-type specificity (**Fig. 5D**). Nevertheless, a subset of SEs was common to red and mosaic s2CLCs, defining a core 2C-like enhancer set (**Fig. 5E**). Fingerprinting analysis of H3K4me3, H3K27ac, and Input confirmed differential coverage across samples (**Fig. S10A–C**).

Together, these results show that s2CLC stability is associated with two layers of enhancer remodeling: (i) ERV-proximal enhancer acetylation that reinforces MERVL activity, and (ii) establishment of cell-type-specific SE landscapes with a shared 2C-like enhancer nucleus.

### Chromatin state modeling reveals ERV integration into H3K27ac-rich states

Finally, we integrated histone modification profiles using ChromHMM^40^. A four-state model identified State 1 (H3K27ac-dominant), State 3 (H3K4me3+H3K27ac), and State 4 (H3K4me3-dominant) as active states (**Fig. 6A**).

**Figure 6:**
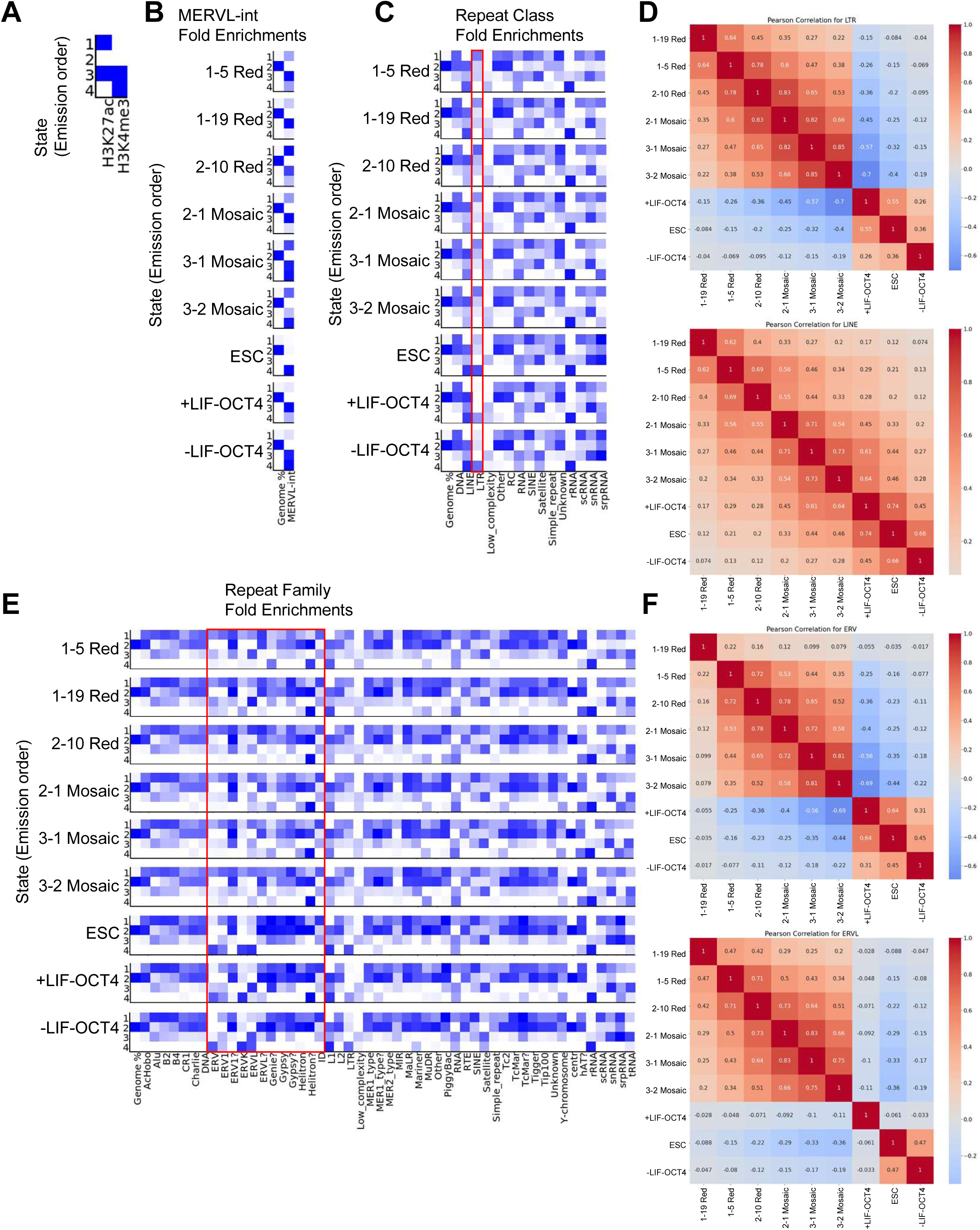
Chromatin state analysis and repeat element enrichment in s2CLCs. (**A**) ChromHMM identification of four chromatin states defined by H3K4me3 and H3K27ac, delineating active regions in s2CLCs. (**B**) ChromHMM enrichment of MERVL-int elements across chromatin states in s2CLCs, conventional ESCs, and ESCs under differentiation conditions. (**C**) ChromHMM enrichment of DNA repeat classes across chromatin states in the same conditions. (**D**) Pearson correlation analysis of ChromHMM enrichment for LTR (top) and LINE (bottom) repeats. (**E**) ChromHMM enrichment of repeat families across chromatin states in ESCs. (**F**) Pearson correlation analysis of ChromHMM enrichment for ERV (top) and ERVL (bottom) repeats.

Neighborhood enrichment revealed distinct distributions of MERVL-int: in ESCs, MERVL-int localized primarily to State 4 (H3K4me3-only), whereas in red s2CLCs it shifted into States 1 and 3 (H3K27ac-enriched), with moderate retention in State 4 (**Fig. 6B**). Mosaic s2CLCs exhibited mixed enrichment across States 1/3/4, while differentiated cells favored States 3 and 4. Thus, red s2CLCs reposition MERVL into H3K27ac-rich states, aligning ERV transcription with enhancer activity.

ChromHMM applied to repeat classes showed that ERV/LTRs were preferentially enriched in red and mosaic s2CLCs compared to ESCs or differentiated cells (**Fig. 6C**). Pearson correlations of ChromHMM enrichments clustered red and mosaic s2CLCs together, apart from ESCs and differentiated cells (**Fig. 6D**). Analysis of RNA repeats supported this separation (**Fig. S11**), with low-complexity and satellite repeats showing distinct profiles. Notably, srpRNA repeats correlated specifically between red s2CLCs and one mosaic clone, pointing to clone-specific features.

At the DNA repeat family level, ERVs localized predominantly to States 1 and 3 in red and mosaic s2CLCs, while conventional and differentiated ESCs favored State 4 (**Fig. 6E**). Other repeat families (ERV1, LTR, RNA repeats) exhibited similar patterns (**Fig. S12**). Pearson correlations further confirmed stronger similarity between red and mosaic s2CLCs than with ESCs or differentiated states (**Fig. 6F**).

These analyses establish that stable 2C-like cells are characterized by the redistribution of ERVs into H3K27ac-rich chromatin states, coupled with enhancer remodeling and SE establishment, which together stabilize the 2C-like transcriptional program.

## DISCUSSION

In this study, we demonstrate that embryonic stem cells (ESCs) can be stably maintained in a 2C-like state, defined by persistent activation of MERVL elements and sustained expression of 2C-specific genes. Unlike the sporadic and heterogeneous bursts of MERVL activity in conventional ESCs, the stable 2C-like cells (s2CLCs) described here exhibit long-term homogeneity, as confirmed by live imaging and flow cytometry. This persistence is accompanied by induction of canonical 2C regulators such as Zscan4 and Dux, transcription factors that are critical for early embryonic transcriptional reprogramming. Dux, a pioneer transcription factor that drives zygotic genome activation (ZGA) and activates MERVL transcription^8, 9^, was strongly upregulated in red s2CLCs. Zscan4, expressed in 2C-stage blastomeres and transiently in ESCs^7^, was also robustly induced. Although only ∼5% of ESCs typically express Zscan4 at any one time, nearly all cells activate it over successive passages^41^. Our results suggest that stabilizing Dux- and Zscan4-positive cells requires an accompanying chromatin environment that permits persistent MERVL activity.

Epigenomic profiling further underscored that s2CLCs possess distinct chromatin states and enhancer architectures, including elevated promoter- and enhancer-associated marks (**Fig. 4**), the redistribution of MERVL elements into H3K27ac-rich domains (**Fig. 6**), and the formation of super enhancer clusters (**Fig. 5**). These features distinguish red and mosaic s2CLCs from conventional ESCs and differentiated states, supporting the idea that stabilization of the 2C-like state involves broad remodeling of promoter and enhancer landscapes. Our analyses of MERVL-associated enhancer acetylation provide a mechanistic link between ERV transcriptional activity and the persistence of the 2C program.

A key finding is the identification of two distinct MERVL-positive subpopulations. Mosaic s2CLCs displayed heterogeneous and fluctuating reporter activity, while red s2CLCs exhibited uniformly high reporter expression and greater stability. These divergent expression patterns were mirrored at the transcriptomic and epigenomic levels: red s2CLCs expressed a broader repertoire of 2C-specific genes and ERVL retrotransposons, whereas mosaic s2CLCs appeared to represent an intermediate or less stable state. This heterogeneity highlights that even within MERVL-positive cells, distinct pathways may govern the acquisition and maintenance of the 2C-like state.

Our repeat expression analyses revealed that red s2CLCs consistently upregulate multiple ERVL family retrotransposons (MERVL-int, MT2_Mm, RLTR6_Mm, ORR1A3-int, GSAT_MM), while mosaic s2CLCs uniquely upregulated RLTR13B2. Our analyses revealed correlative upregulation of RLTR6_Mm, ORR1A3-int, and GSAT_MM in red s2CLCs, suggesting these elements may serve as additional markers of the stable 2C-like state. Prior studies have shown that MERVL elements function as alternative promoters for early embryonic genes and modulate chromatin accessibility^4, 42, 43^. Our data build on these findings by showing that different ERVL subfamilies contribute variably to 2C identity depending on cellular context, suggesting specialized regulatory roles for distinct retrotransposons in stabilizing or destabilizing totipotency.

The connection between ERV activity, transcription factor networks, and chromatin remodeling revealed here provides new insight into how ESCs toggle between pluripotency and totipotency. Evolutionary co-option of retrotransposons into host regulatory networks may explain why MERVL and related ERVs are central to ZGA and to the 2C program. By capturing a stable 2C-like state, we provide a tractable in vitro system to interrogate how ERV-driven transcription shapes developmental potential and to test whether modulation of enhancer states can shift cell fate.

Functionally, s2CLCs demonstrated altered differentiation outcomes in embryoid body assays, including increased morphological heterogeneity and the emergence of cystic structures associated with trophectoderm potential. These results support the idea that stabilization of the 2C program directly influences lineage trajectories, reinforcing the link between ERV activation, chromatin remodeling, and expanded developmental potency. Conventional ESC cultures, in which MERVL+ cells are rare and transient, are limited for such studies. By contrast, s2CLCs provide an enriched and reproducible model for dissecting totipotent-like features.

Taken together, our findings establish that the stable maintenance of 2C-like cells is achievable in vitro and that this state is defined by coordinated regulation of transcription factors, retrotransposons, and chromatin states. By elucidating the molecular and epigenetic features of s2CLCs, our study extends current understanding of how ESCs access the 2C-like state and demonstrates that this state can be captured and sustained. These results not only illuminate fundamental principles of early embryonic gene regulation but also provide a foundation for future investigations into ERV biology, reprogramming, and regenerative medicine. In summary, the establishment of stable 2C-like cells offers a robust new model for investigating the interplay between retroelements, chromatin organization, and developmental potential in mammalian cells.

## MATERIALS & METHODS

### ESC cell culture

ESCs were cultured according to previously established methods with some alterations^44, 45^. For this study, we used an embryonic stem cell line that allows conditional modulation of OCT4 levels^27^. These cells are widely employed to study both self-renewal and differentiation, as they retain pluripotency under maintenance conditions yet can be driven to differentiate through OCT4 downregulation. ESCs were maintained on gelatin-coated dishes in ESC medium composed of DMEM supplemented with 15% FBS and LIF (ESGRO) at 37 °C with 5% CO₂, with medium additionally supplemented with 1.5 µM CHIR9901 (a GSK3 inhibitor). Under these conditions (LIF present, doxycycline absent), cells are referred to as conventional ESCs. For OCT4 downregulation, ESCs were cultured in the same medium supplemented with 2 µg/mL doxycycline. For differentiation, ESCs were cultured without LIF, either with or without doxycycline. Cells were routinely passaged after PBS washing and trypsin dissociation using serological pipettes (sc-200279, sc-200281).

### Isolation of MERVL-positive reporter ESCs

A 2C::tdTomato reporter plasmid^4^ was used. Modifications were made to this plasmid to incorporate a selectable cassette. ESCs were transfected with the modified reporter plasmid using Lipofectamine 2000. Following transfection, cells were selected and subsequently expanded. ESCs were used to generate stable MERVL promoter-driven tdTomato^4^ integrations (**Fig. 1A**).

### Flow cytometry analysis

ESCs were washed with 1X PBS, trypsinized to a single-cell suspension, and re-suspended at a density of 10^6^ cells/mL in FACS buffer (PBS + 1% FBS). Cells were sorted using gates set to include 2C::tdTomato positive control ESCs. A BD LSR II instrument was used for flow cytometry analysis. Data were analyzed using FloJo (FlowJo, LLC). Three biological replicates were performed.

### Real-time imaging

The 2C::tdTomato ES cells were imaged every 30 minutes over a period of 72 hours using the Sartorius IncuCyte SX 5 live-cell imaging system. The analysis focused on assessing the tdTomato area relative to the phase area (which corresponds to the ES cell colony size) and the total tdTomato area, measured in µm^2^/image. Representative images are displayed in **Figure 1I-J**.

### ChIP-Seq analysis

ChIP-Seq experiments followed previously described methods with some modifications^46,47^. We used the rabbit monoclonal antibody H3K4me3 (17-614) from Millipore and the rabbit polyclonal H3K27ac (ab4729) from Abcam. Briefly, ESCs were crosslinked using 1% formaldehyde for 10 minutes at 37°C. The fixed cells were then frozen at −80°C. Later, these cell pellets were thawed and sonicated. For ChIP assays, extracts from 4 million cells were used along with 4 µg of antibody. The ChIP-enriched DNA was processed using the NEBNext Ultra II End Repair/dA-Tailing module, followed by ligation of Illumina adapters. PCR was conducted with the Phusion 2X High Fidelity PCR master mix. The prepared ChIP libraries were then sequenced using Illumina HiSeq platforms following the manufacturer’s protocols. Sequence reads were aligned to the mouse genome via bowtie2^48^ using default settings. For data processing, previously outlined C++ programs^49^ were utilized for: converting SAM-formatted files to BED6 format from bowtie2 (Sam2Bed6_Bowtie2), removing redundant reads from a BED6 file (RemoveRedundantReads), and converting a BED6 file to a BEDGraph file (GenerateRPBMBasedSummary).

ChIP-Seq enriched regions (peaks) were detected by comparing them against control Input DNA using SICER^36^. The settings applied were a window size of 200 bp, a gap size of 400 bp, and an FDR of 0.001. For analyzing multiple samples, the SICER-compare function was used with criteria of FDR < 0.001 and a fold-change > 1.5. Normalization of ChIP-Seq libraries was carried out based on the size of the library, employing the RPBM metric (reads per base per million reads) to measure densities in genomic areas from ChIP-Seq datasets. The analysis included two biological replicates. Visualization of the normalized ChIP data was achieved through the UCSC genome browser.

### RNA-Seq analysis

Poly-A mRNA was extracted using the NEBNext Ultra II RNA Library Prep Kit for Illumina. RNA-Seq libraries were sequenced on an Illumina system in accordance with the manufacturer’s instructions. Sequence reads were aligned to the mouse genome using bowtie2^48^ with default settings. The RPKM metric (reads per kilobase of exon model per million reads)^50^ was used to measure the mRNA expression levels of genes from RNA-Seq data. Genes showing differential expression were determined using edgeR, applying criteria of an FDR less than 0.001 and a fold change greater than 1.5)^51^. RNA-Seq analysis was conducted with three biological replicates.

### Chromatin state learning

Chromatin states in MERVL-positive ESCs were determined using ChromHMM v1.2^52^, which utilizes a multivariate Hidden Markov Model approach. The ChromHMM model was configured by merging data on histone modifications (ChIP-enriched peaks, refer to ChIP-Seq methods described previously) specifically for H3K4me3 and H3K27ac. For each ChIP-Seq dataset, peaks were evaluated in 200 bp bin intervals across the genome. These bins were classified into two categories: 1 for peak enrichment and 0 for no enrichment. A four-state model was chosen because it effectively identified key combinatorial patterns of histone modifications across the genome.

### Chromatin state annotations

The four chromatin states in the ESC epigenome were annotated using CpG islands from the UCSC Genome Browser, and genic features such as transcription start sites (TSS), transcription end sites (TES), genes, exons, and introns were incorporated into ChromHMM using ENCODE annotations, following previously described methods.

### Super enhancer analysis

Super enhancers marked by H3K27ac were detected using HOMER^38^. Initially, all enhancers were evaluated using the findPeaks function of HOMER and plotted with the addition of the “-superSlope −1000” parameter. H3K27ac peaks within 12.5 kb of each other were consolidated. The signal of each super enhancer area was derived by deducting the count of normalized input reads from the count of normalized reads. These areas were then sorted, scaled relative to the highest value, and the quantity of typical enhancer regions was calculated. Super enhancers were designated as areas exceeding a slope of 1 (slope >1).

### Metascape network analysis

Metascape^29^ was used to perform gene ontology (GO) functional annotation and to provide functional insights into genes. This included GO enrichment analysis covering biological processes, canonical pathways, and WikiPathways results. Additionally, protein network analyses were carried out using Metascape.

### Heatmaps

The “computeMatrix” and “plotHeatmap” functions in the DeepTools^39^ software suite were used to generate heatmaps and profile plots for histone marks and transcription factor binding from ChIP-seq datasets. These visualizations focused on regions extending from −3kb to +3kb around the peak centers.

## DECLARATIONS

### ETHICS APPROVAL AND CONSENT TO PARTICIPATE

Not applicable

### CONSENT FOR PUBLICATION

All authors have read and approved the final version of this manuscript.

## Data Availability

The sequencing data from this study have been submitted to the NCBI Gene Expression Omnibus (GEO).

## Statistics and Reproducibility

We generated biological duplicate H3K4me3 and H3K27ac ChIP-Seq datasets for teratomas. We also generated biological triplicate RNA-Seq datasets.

## AUTHORS’ CONTRIBUTIONS

B.L.K. conceived of the study, designed and carried out the experiments, analyzed the sequencing data, and drafted the manuscript. R.G. assisted with experiments such as EB differentiation assays, and NGS library preparation.

## ACKNOWLEDGEMENTS

This work utilized the Wayne State University High Performance Computing Grid for computational resources (https://www.grid.wayne.edu/). Flow cytometry was performed in the Microscopy, Imaging, and Cytometry Resources (MICR) core at the Karmanos Cancer Institute and Wayne State University.

## REFERENCES

1. Tarkowski AK. Experiments on the development of isolated blastomeres of mouse eggs. Nature. 1959;184(4695):1286–7.

2. Wu G, Scholer HR. Lineage Segregation in the Totipotent Embryo. Curr Top Dev Biol. 2016;117:301–17. doi: 10.1016/bs.ctdb.2015.10.014. PubMed PMID: 26969985.

3. Borsos M, Torres-Padilla M-E. Building up the nucleus: nuclear organization in the establishment of totipotency and pluripotency during mammalian development. Genes & development. 2016;30(6):611–21.

4. Macfarlan TS, Gifford WD, Driscoll S, Lettieri K, Rowe HM, Bonanomi D, Firth A, Singer O, Trono D, Pfaff SL. Embryonic stem cell potency fluctuates with endogenous retrovirus activity. Nature. 2012;487(7405):57–63. Epub 2012/06/23. doi: nature11244 [pii] 10.1038/nature11244. PubMed PMID: 22722858; PMCID: 3395470.

5. Fadloun A, Le Gras S, Jost B, Ziegler-Birling C, Takahashi H, Gorab E, Carninci P, Torres-Padilla ME. Chromatin signatures and retrotransposon profiling in mouse embryos reveal regulation of LINE-1 by RNA. Nature structural & molecular biology. 2013;20(3):332–8. doi: 10.1038/nsmb.2495. PubMed PMID: 23353788.

6. Peaston AE, Evsikov AV, Graber JH, de Vries WN, Holbrook AE, Solter D, Knowles BB. Retrotransposons regulate host genes in mouse oocytes and preimplantation embryos. Dev Cell. 2004;7(4):597–606. Epub 2004/10/08. doi: S1534580704003259 [pii] 10.1016/j.devcel.2004.09.004. PubMed PMID: 15469847.

7. Falco G, Lee SL, Stanghellini I, Bassey UC, Hamatani T, Ko MS. Zscan4: a novel gene expressed exclusively in late 2-cell embryos and embryonic stem cells. Dev Biol. 2007;307(2):539–50. doi: 10.1016/j.ydbio.2007.05.003. PubMed PMID: 17553482; PMCID: PMC1994725.

8. De Iaco A, Planet E, Coluccio A, Verp S, Duc J, Trono D. DUX-family transcription factors regulate zygotic genome activation in placental mammals. Nat Genet. 2017;49(6):941–5. doi: 10.1038/ng.3858. PubMed PMID: 28459456; PMCID: PMC5446900.

9. Hendrickson PG, Dorais JA, Grow EJ, Whiddon JL, Lim JW, Wike CL, Weaver BD, Pflueger C, Emery BR, Wilcox AL, Nix DA, Peterson CM, Tapscott SJ, Carrell DT, Cairns BR. Conserved roles of mouse DUX and human DUX4 in activating cleavage-stage genes and MERVL/HERVL retrotransposons. Nat Genet. 2017;49(6):925–34. doi: 10.1038/ng.3844. PubMed PMID: 28459457; PMCID: PMC5703070.

10. Evsikov AV, de Vries WN, Peaston AE, Radford EE, Fancher KS, Chen FH, Blake JA, Bult CJ, Latham KE, Solter D, Knowles BB. Systems biology of the 2-cell mouse embryo. Cytogenet Genome Res. 2004;105(2-4):240–50. doi: 10.1159/000078195. PubMed PMID: 15237213.

11. Kigami D, Minami N, Takayama H, Imai H. MuERV-L is one of the earliest transcribed genes in mouse one-cell embryos. Biol Reprod. 2003;68(2):651–4. doi: 10.1095/biolreprod.102.007906. PubMed PMID: 12533431.

12. Svoboda P, Stein P, Anger M, Bernstein E, Hannon GJ, Schultz RM. RNAi and expression of retrotransposons MuERV-L and IAP in preimplantation mouse embryos. Dev Biol. 2004;269(1):276–85. doi: 10.1016/j.ydbio.2004.01.028. PubMed PMID: 15081373.

13. Ribet D, Louvet-Vallee S, Harper F, de Parseval N, Dewannieux M, Heidmann O, Pierron G, Maro B, Heidmann T. Murine endogenous retrovirus MuERV-L is the progenitor of the “orphan” epsilon viruslike particles of the early mouse embryo. J Virol. 2008;82(3):1622–5. doi: 10.1128/JVI.02097-07. PubMed PMID: 18045933; PMCID: PMC2224431.

14. Friedli M, Trono D. The developmental control of transposable elements and the evolution of higher species. Annual review of cell and developmental biology. 2015;31:429–51.

15. De Los Angeles A, Ferrari F, Xi R, Fujiwara Y, Benvenisty N, Deng H, Hochedlinger K, Jaenisch R, Lee S, Leitch HG. Hallmarks of pluripotency. Nature. 2015;525(7570):469–78.

16. Lee HJ, Hore TA, Reik W. Reprogramming the methylome: erasing memory and creating diversity. Cell stem cell. 2014;14(6):710–9.

17. Torres-Padilla M-E, Chambers I. Transcription factor heterogeneity in pluripotent stem cells: a stochastic advantage. Development. 2014;141(11):2173–81.

18. Ishiuchi T, Enriquez-Gasca R, Mizutani E, Bošković A, Ziegler-Birling C, Rodriguez-Terrones D, Wakayama T, Vaquerizas JM, Torres-Padilla M-E. Early embryonic-like cells are induced by downregulating replication-dependent chromatin assembly. Nature structural & molecular biology. 2015;22(9):662–71.

19. Dan J, Yang J, Liu Y, Xiao A, Liu L. Roles for histone acetylation in regulation of telomere elongation and two-cell state in mouse ES cells. Journal of cellular physiology. 2015;230(10):2337–44.

20. Walter M, Teissandier A, Pérez-Palacios R, Bourc’His D. An epigenetic switch ensures transposon repression upon dynamic loss of DNA methylation in embryonic stem cells. Elife. 2016;5:e11418.

21. Hayashi K, Lopes SM, Tang F, Surani MA. Dynamic equilibrium and heterogeneity of mouse pluripotent stem cells with distinct functional and epigenetic states. Cell Stem Cell. 2008;3(4):391–401. doi: 10.1016/j.stem.2008.07.027. PubMed PMID: 18940731; PMCID: PMC3847852.

22. Thompson PJ, Dulberg V, Moon K-M, Foster LJ, Chen C, Karimi MM, Lorincz MC. hnRNP K coordinates transcriptional silencing by SETDB1 in embryonic stem cells. PLoS genetics. 2015;11(1):e1004933.

23. Hisada K, Sánchez C, Endo TA, Endoh M, Román-Trufero M, Sharif J, Koseki H, Vidal M. RYBP represses endogenous retroviruses and preimplantation-and germ line-specific genes in mouse embryonic stem cells. Molecular and cellular biology. 2012;32(6):1139–49.

24. Macfarlan TS, Gifford WD, Agarwal S, Driscoll S, Lettieri K, Wang J, Andrews SE, Franco L, Rosenfeld MG, Ren B. Endogenous retroviruses and neighboring genes are coordinately repressed by LSD1/KDM1A. Genes & development. 2011;25(6):594–607.

25. Maksakova IA, Thompson PJ, Goyal P, Jones SJ, Singh PB, Karimi MM, Lorincz MC. Distinct roles of KAP1, HP1 and G9a/GLP in silencing of the two-cell-specific retrotransposon MERVL in mouse ES cells. Epigenetics Chromatin. 2013;6(1):15. doi: 10.1186/1756-8935-6-15. PubMed PMID: 23735015; PMCID: PMC3682905.

26. Rowe HM, Jakobsson J, Mesnard D, Rougemont J, Reynard S, Aktas T, Maillard PV, Layard-Liesching H, Verp S, Marquis J, Spitz F, Constam DB, Trono D. KAP1 controls endogenous retroviruses in embryonic stem cells. Nature. 2010;463(7278):237–40. Epub 2010/01/16. doi: nature08674 [pii] 10.1038/nature08674. PubMed PMID: 20075919.

27. Niwa H, Miyazaki J, Smith AG. Quantitative expression of Oct-3/4 defines differentiation, dedifferentiation or self-renewal of ES cells. Nat Genet. 2000;24(4):372–6. PubMed PMID: 10742100.

28. Yu G, Wang LG, Han Y, He QY. clusterProfiler: an R package for comparing biological themes among gene clusters. Omics : a journal of integrative biology. 2012;16(5):284–7. Epub 2012/03/30. doi: 10.1089/omi.2011.0118. PubMed PMID: 22455463; PMCID: PMC3339379.

29. Zhou Y, Zhou B, Pache L, Chang M, Khodabakhshi AH, Tanaseichuk O, Benner C, Chanda SK. Metascape provides a biologist-oriented resource for the analysis of systems-level datasets. Nat Commun. 2019;10(1):1523. doi: 10.1038/s41467-019-09234-6. PubMed PMID: 30944313; PMCID: PMC6447622.

30. Shannon P, Markiel A, Ozier O, Baliga NS, Wang JT, Ramage D, Amin N, Schwikowski B, Ideker T. Cytoscape: a software environment for integrated models of biomolecular interaction networks. Genome Res. 2003;13(11):2498–504. doi: 10.1101/gr.1239303. PubMed PMID: 14597658; PMCID: PMC403769.

31. Barski A, Cuddapah S, Cui K, Roh TY, Schones DE, Wang Z, Wei G, Chepelev I, Zhao K. High-resolution profiling of histone methylations in the human genome. Cell. 2007;129(4):823–37. Epub 2007/05/22. doi: S0092-8674(07)00600-9 [pii] 10.1016/j.cell.2007.05.009. PubMed PMID: 17512414.

32. Sims RJ, 3rd, Nishioka K, Reinberg D. Histone lysine methylation: a signature for chromatin function. Trends Genet. 2003;19(11):629–39. Epub 2003/10/31. doi: S0168-9525(03)00261-0 [pii] 10.1016/j.tig.2003.09.007. PubMed PMID: 14585615.

33. Santos-Rosa H, Schneider R, Bannister AJ, Sherriff J, Bernstein BE, Emre NC, Schreiber SL, Mellor J, Kouzarides T. Active genes are tri-methylated at K4 of histone H3. Nature. 2002;419(6905):407–11. Epub 2002/09/28. doi: 10.1038/nature01080 nature01080 [pii]. PubMed PMID: 12353038.

34. Schneider R, Bannister AJ, Myers FA, Thorne AW, Crane-Robinson C, Kouzarides T. Histone H3 lysine 4 methylation patterns in higher eukaryotic genes. Nat Cell Biol. 2004;6(1):73–7. Epub 2003/12/09. doi: 10.1038/ncb1076 ncb1076 [pii]. PubMed PMID: 14661024.

35. Whyte WA, Orlando DA, Hnisz D, Abraham BJ, Lin CY, Kagey MH, Rahl PB, Lee TI, Young RA. Master transcription factors and mediator establish super-enhancers at key cell identity genes. Cell. 2013;153(2):307–19. doi: 10.1016/j.cell.2013.03.035. PubMed PMID: 23582322; PMCID: PMC3653129.

36. Zang C, Schones DE, Zeng C, Cui K, Zhao K, Peng W. A clustering approach for identification of enriched domains from histone modification ChIP-Seq data. Bioinformatics. 2009;25(15):1952–8. Epub 2009/06/10. doi: btp340 [pii] 10.1093/bioinformatics/btp340. PubMed PMID: 19505939.

37. Khan A, Mathelier A. Intervene: a tool for intersection and visualization of multiple gene or genomic region sets. BMC Bioinformatics. 2017;18(1):287. doi: 10.1186/s12859-017-1708-7. PubMed PMID: 28569135; PMCID: PMC5452382.

38. Heinz S, Benner C, Spann N, Bertolino E, Lin YC, Laslo P, Cheng JX, Murre C, Singh H, Glass CK. Simple combinations of lineage-determining transcription factors prime cis-regulatory elements required for macrophage and B cell identities. Mol Cell. 2010;38(4):576–89. doi: 10.1016/j.molcel.2010.05.004. PubMed PMID: 20513432; PMCID: 2898526.

39. Ramirez F, Dundar F, Diehl S, Gruning BA, Manke T. deepTools: a flexible platform for exploring deep-sequencing data. Nucleic Acids Res. 2014;42(Web Server issue):W187-91. doi: 10.1093/nar/gku365. PubMed PMID: 24799436; PMCID: PMC4086134.

40. Ernst J, Kellis M. Chromatin-state discovery and genome annotation with ChromHMM. Nat Protoc. 2017;12(12):2478–92. doi: 10.1038/nprot.2017.124. PubMed PMID: 29120462; PMCID: PMC5945550.

41. Zalzman M, Falco G, Sharova LV, Nishiyama A, Thomas M, Lee SL, Stagg CA, Hoang HG, Yang HT, Indig FE, Wersto RP, Ko MS. Zscan4 regulates telomere elongation and genomic stability in ES cells. Nature. 2010;464(7290):858–63. doi: 10.1038/nature08882. PubMed PMID: 20336070; PMCID: PMC2851843.

42. Eckersley-Maslin MA, Svensson V, Krueger C, Stubbs TM, Giehr P, Krueger F, Miragaia RJ, Kyriakopoulos C, Berrens RV, Milagre I. MERVL/Zscan4 network activation results in transient genome-wide DNA demethylation of mESCs. Cell reports. 2016;17(1):179–92.

43. Iturbide A, Torres-Padilla ME. A cell in hand is worth two in the embryo: recent advances in 2-cell like cell reprogramming. Curr Opin Genet Dev. 2020;64:26–30. Epub 2020/07/01. doi: 10.1016/j.gde.2020.05.038. PubMed PMID: 32599301.

44. Kidder BL, Hu G, Yu ZX, Liu C, Zhao K. Extended self-renewal and accelerated reprogramming in the absence of Kdm5b. Mol Cell Biol. 2013;33(24):4793–810. Epub 2013/10/09. doi: MCB.00692-13 [pii] 10.1128/MCB.00692-13. PubMed PMID: 24100015.

45. Kidder BL, Hu G, Zhao K. KDM5B focuses H3K4 methylation near promoters and enhancers during embryonic stem cell self-renewal and differentiation. Genome Biol. 2014;15(2):R32. Epub 2014/02/06. doi: gb-2014-15-2-r32 [pii] 10.1186/gb-2014-15-2-r32. PubMed PMID: 24495580.

46. Gopi LK, Kidder BL. Integrative pan cancer analysis reveals epigenomic variation in cancer type and cell specific chromatin domains. Nat Commun. 2021;12(1):1419. doi: 10.1038/s41467-021-21707-1. PubMed PMID: 33658503; PMCID: PMC7930052.

47. He R, Xhabija B, Gopi LK, Kurup JT, Xu Z, Liu Z, Kidder BL. H3K4 demethylase KDM5B regulates cancer cell identity and epigenetic plasticity. Oncogene. 2022;41(21):2958–72. doi: 10.1038/s41388-022-02311-z. PubMed PMID: 35440714; PMCID: PMC9426628.

48. Langmead B, Salzberg SL. Fast gapped-read alignment with Bowtie 2. Nat Methods. 2012;9(4):357–9. Epub 2012/03/06. doi: nmeth.1923 [pii] 10.1038/nmeth.1923. PubMed PMID: 22388286; PMCID: 3322381.

49. Hu G, Zhao K. Correlating histone modification patterns with gene expression data during hematopoiesis. Methods Mol Biol. 2014;1150:175–87. doi: 10.1007/978-1-4939-0512-6_11. PubMed PMID: 24743998; PMCID: PMC4198375.

50. Mortazavi A, Williams BA, McCue K, Schaeffer L, Wold B. Mapping and quantifying mammalian transcriptomes by RNA-Seq. Nat Methods. 2008;5(7):621–8. Epub 2008/06/03. doi: nmeth.1226 [pii] 10.1038/nmeth.1226. PubMed PMID: 18516045.

51. Robinson MD, McCarthy DJ, Smyth GK. edgeR: a Bioconductor package for differential expression analysis of digital gene expression data. Bioinformatics. 2009;26(1):139–40. Epub 2009/11/17. doi: btp616 [pii] 10.1093/bioinformatics/btp616. PubMed PMID: 19910308.

52. Ernst J, Kellis M. ChromHMM: automating chromatin-state discovery and characterization. Nat Methods. 2012;9(3):215–6. doi: 10.1038/nmeth.1906. PubMed PMID: 22373907; PMCID: PMC3577932.

