## Supplemental Figure for "Stable Maintenance of Two-Cell-Like Cells from Embryonic Stem Cells Reveals Chromatin and Super Enhancer Regulation of MERVL Elements"

Running title: Stable maintenance of MERVL-positive ESCs.

[*]Correspondence:

Benjamin L. Kidder

### Supplemental Figures

**Figure S1. Flow cytometry analysis of MERVL expression in ESCs.**

Flow cytometry analysis of the 2C::tdTomato reporter to detect MERVL expression in ESCs.

**Figure S2. Differential RNA-Seq expression analysis of s2CLCs.**

Volcano plot of differentially expressed genes identified between red and mosaic s2CLCs and conventional ESCs. Points represent genes, plotted by log2 fold change (log2FC) on the x-axis and p-value significance on the y-axis. Grey points are not significantly differentially expressed (NS), green points signify significant log2FC, blue points indicate significant p-values, and red points mark genes with both significant log2FC and p-values, highlighting the most differentially expressed genes in red and mosaic s2CLCs vs. conventional ESC comparison.

**Figure S3. MA plots of differential expression in s2CLCs relative to conventional ESCs.**

MA-plots were used to compare gene expression in red and mosaic s2CLCs to that in conventional ESCs. The plot maps the normalized mean expression levels on the x-axis against the log2 fold changes on the y-axis for each gene. Red dots in the plot denote genes that are significantly downregulated, whereas blue dots represent those that are upregulated.

**Figure S4. Gene ontology analysis of differentially expressed genes in s2CLCs.**

Clusterprofiler GO term analysis of genes that are upregulated (left; activated) or downregulated (right; suppressed) in red (left panels) and mosaic (right panels) s2CLCs compared to conventional controls. Enriched GO terms highlight the biological processes and pathways predominantly activated or repressed in these distinct ESC populations, revealing key pathways that are differentially regulated between the cell types.

**Figure S5. TRRUST analysis of transcription factor targets in s2CLCs using metascape.**

TRRUST analysis identifies and highlights the targets of transcription factors (TFs) whose expression is notably enriched in red s2CLCs.

**Figure S6. Volcano plots and heat map clustering of differentially expressed genes in s2CLCs.**

(**A**) Volcano plots display the differential expression of repeat elements (left) and family members (right) between red and mosaic s2CLCs compared to conventional ESCs. Each point represents a gene, plotted by log2 fold change (log2FC) on the x-axis and p-value significance on the y-axis. Grey points indicate genes without significant differential expression, green points denote significant log2FC, blue points represent significant p-values, and red points highlight genes that are significantly different in both log2FC and p-values, showcasing the most distinctively expressed genes in the comparison between red and mosaic s2CLCs versus conventional ESCs. (**B**) and (**C**) Heat maps illustrate the clustering of differential expression for repeat elements (**B**) and family members (**C**) in red and mosaic s2CLCs relative to conventional ESCs, highlighting patterns of gene expression variations across these groups.

**Figure S7. Day 12 embryoid body differentiation and morphological segmentation**

(**A**) Tiled image showing automated segmentation of embryoid bodies, with each color denoting a distinct EB. (**B**) Bright field microscopy tiled image displaying the differentiation of EBs at day 12, highlighting red and mosaic s2CLCs.

**Figure S8. Enrichment of gene ontology (GO) terms in differentiated expressed genes of red s2CLCs during EB differentiation compared to conventional ESCs**

Clusterprofiler analysis reveals GO terms enriched in both activated and suppressed genes for groups of 1-5 red s2CLCs (left) and 1-19 red s2CLCs (right), compared to conventional EBs.

**Figure S9. Epigenome profiling of H3K4me3 and H3K27ac in s2CLCs.**

HOMER annotation of (**A**) H3K4me3 and (**B**) H3K27ac peaks in red and mosaic s2CLCs, conventional ESCs, and ESCs cultured in differentiation conditions, which revealed enrichment of peaks in promoter, intergenic, and intronic regions. deepTools was used to generate heatmap density profiles of (**C**) H3K4me3 and (**D**) H3K27ac ChIP-Seq signals in ESCs around transcriptional start sites (TSS) and gene body regions generated. Row linked heatmaps show k-means clusters of genes with similar histone modification profiles. Scatter plot of (**E**) H3K4me3 and (**F**) H3K27ac densities in red and mosaic s2CLCs relative to conventional ESCs.

**Figure S10. Fingerprints of ChIP signal-to-noise ratios.**

Deeptools facilitated the generation of ChIP-seq fingerprint profiles, which illustrate the cumulative sum of per-base coverage across each analyzed genomic bin.

**Figure S11. Pearson correlation of ChromHMM enrichment for DNA repeat across chromatin states in s2CLCs.**

Pearson correlation analysis of ChromHMM enrichment analysis of repeat classes in different chromatin states identified in s2CLCs, ESCs, and differentiated cells. The analysis provides insights into the relationships and patterns of chromatin organization associated with repeat elements in these cells.

**Figure S12. Pearson correlation of ChromHMM enrichment for DNA family members across chromatin states in ESCs.**

Pearson correlation analysis of ChromHMM enrichment analysis of repeat family members in different chromatin states identified in s2CLCs, ESCs, and differentiated cells.

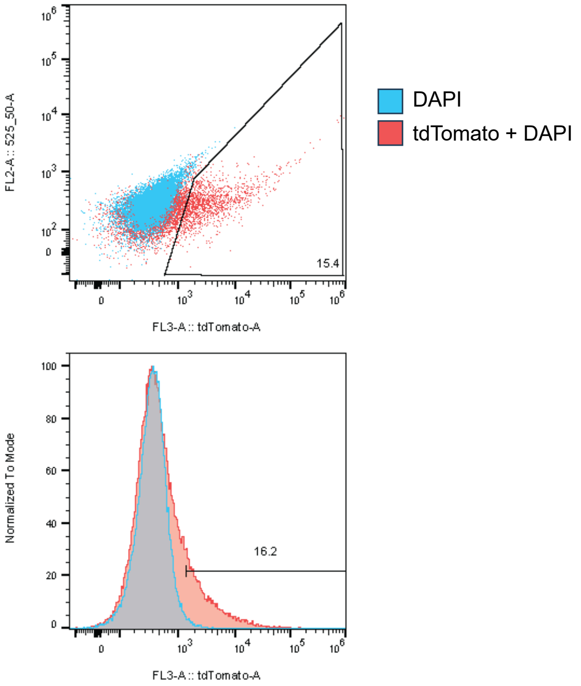
DAPI
tdTomato + DAPI
FL2-A :: 525_50-A
FL3-A :: tdTomato-A
15.4
Normalized To Mode
16.2
FL3-A :: tdTomato-A

Figure S1

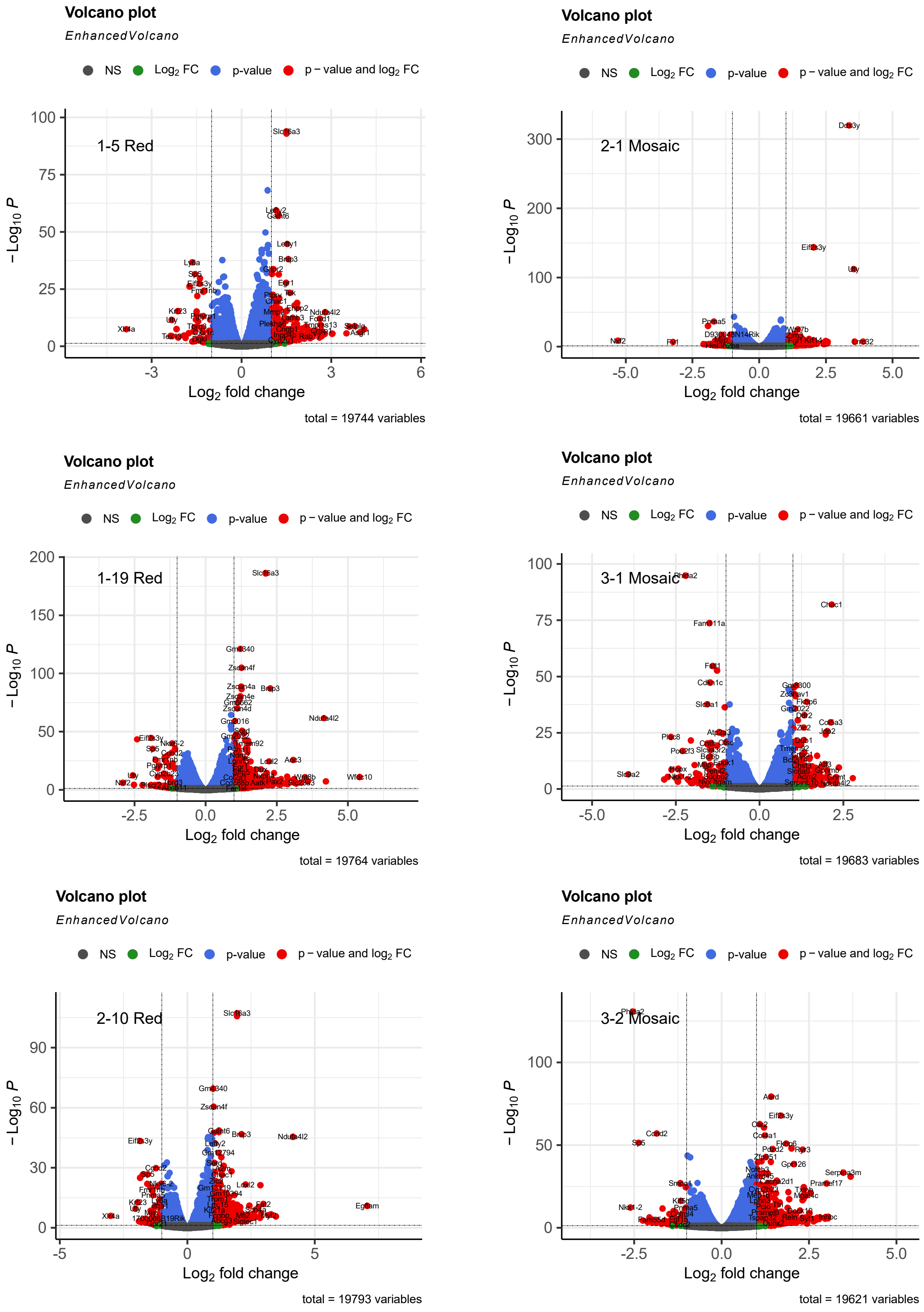

Volcano plot
EnhancedVolcano
NS
Log2 FC
p-value
p – value and log2 FC
1-5 Red
Slc16a3
– Log10 P
Log2 fold change
total = 19744 variables
Volcano plot
EnhancedVolcano
NS
Log2 FC
p-value
p – value and log2 FC
2-1 Mosaic
Ddx3y
Eif2s3y
Uty
– Log10 P
Log2 fold change
total = 19661 variables
Volcano plot
EnhancedVolcano
NS
Log2 FC
p-value
p – value and log2 FC
1-19 Red
Slc16a3
Zscan4f
Brip3
Ndufa4l2
Wfdc10
– Log10 P
Log2 fold change
total = 19764 variables
Volcano plot
EnhancedVolcano
NS
Log2 FC
p-value
p – value and log2 FC
3-1 Mosaic
Chac1
Fam111a
Slc6a2
– Log10 P
Log2 fold change
total = 19683 variables
Volcano plot
EnhancedVolcano
NS
Log2 FC
p-value
p – value and log2 FC
2-10 Red
Slc16a3
Zscan4f
Eif2s3y
Ndufa4l2
Egfem
– Log10 P
Log2 fold change
total = 19793 variables
Volcano plot
EnhancedVolcano
NS
Log2 FC
p-value
p – value and log2 FC
3-2 Mosaic
Eif2s3y
Serpina3m
Pramef17
– Log10 P
Log2 fold change
total = 19621 variables


Figure S2

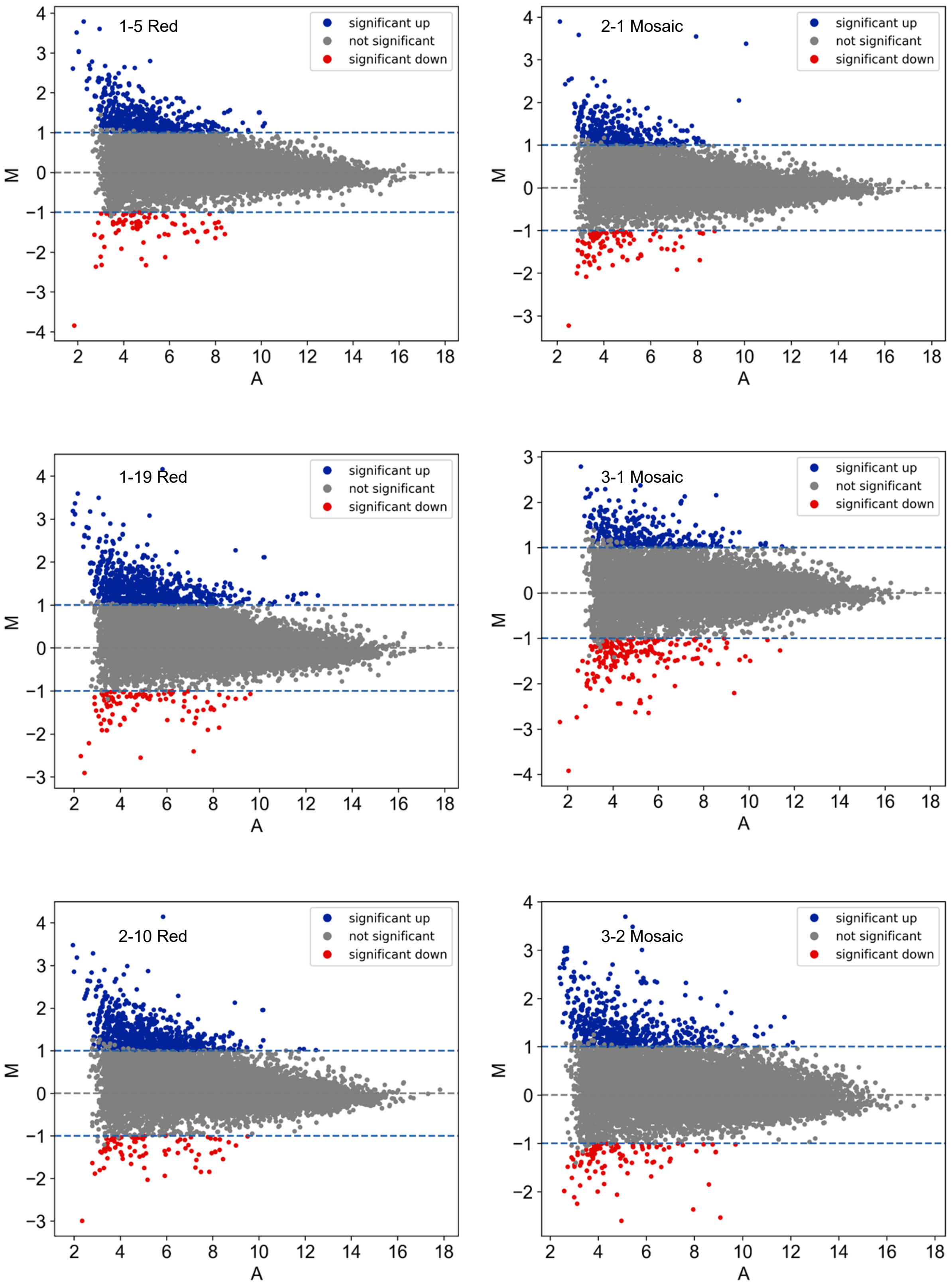
1-5 Red
significant up
not significant
significant down
M
A
2-1 Mosaic
significant up
not significant
significant down
M
A
1-19 Red
significant up
not significant
significant down
M
A
3-1 Mosaic
significant up
not significant
significant down
M
A
2-10 Red
significant up
not significant
significant down
M
A
3-2 Mosaic
significant up
not significant
significant down
M
A

Figure S3

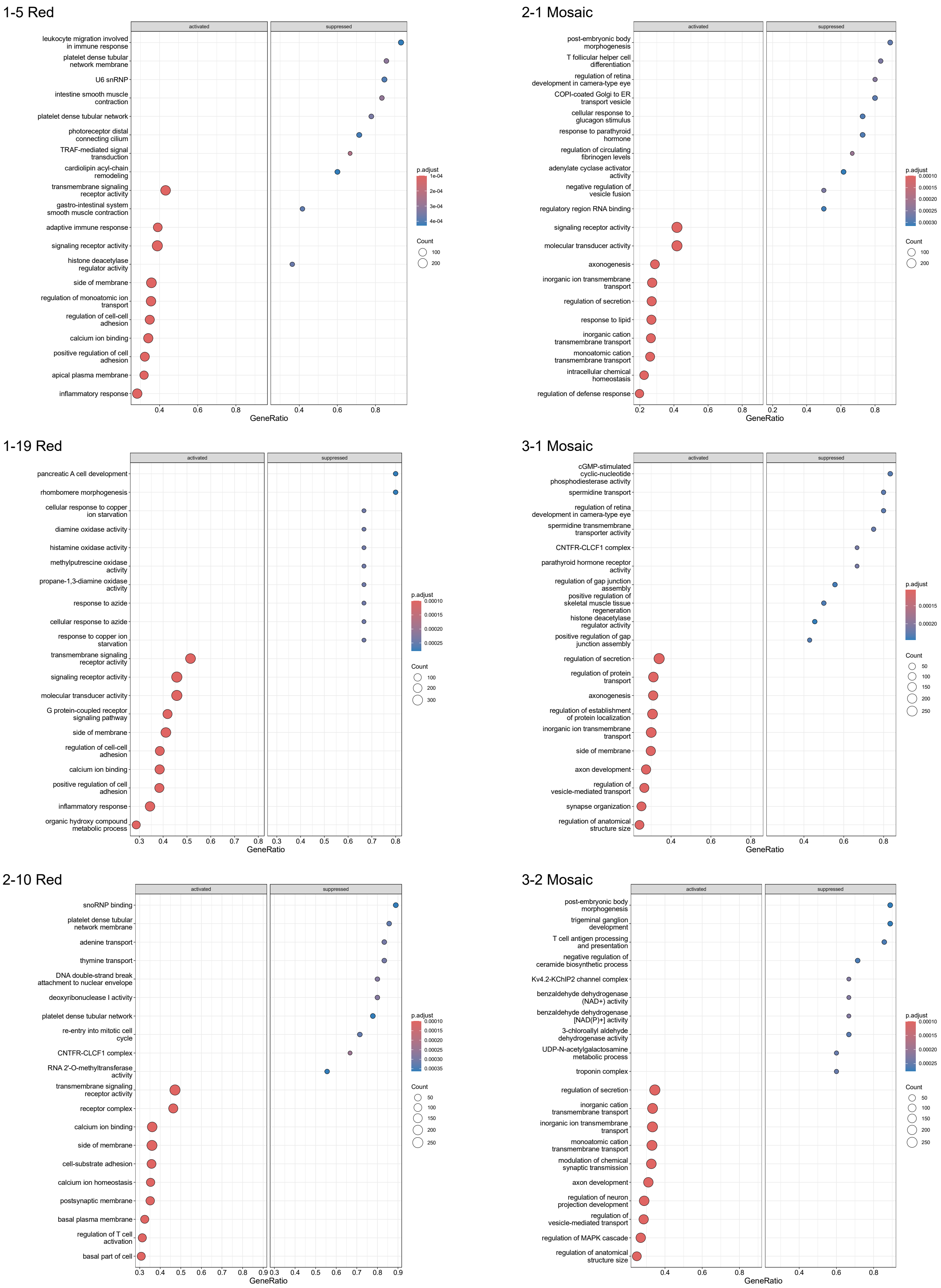
1-5 Red
activated
suppressed
leukocyte migration involved in immune response
platelet dense tubular network membrane
U6 snRNP
intestine smooth muscle contraction
platelet dense tubular network
photoreceptor distal connecting cilium
TRAF-mediated signal transduction
cardiolipin acyl-chain remodeling
transmembrane signaling receptor activity
gastro-intestinal system smooth muscle contraction
adaptive immune response
signaling receptor activity
histone deacetylase regulator activity
side of membrane
regulation of monoatomic ion transport
regulation of cell-cell adhesion
calcium ion binding
positive regulation of cell adhesion
apical plasma membrane
inflammatory response
GeneRatio
p.adjust
Count
2-1 Mosaic
post-embryonic body morphogenesis
T follicular helper cell differentiation
regulation of retina development in camera-type eye
COPI-coated Golgi to ER transport vesicle
cellular response to glucagon stimulus
response to parathyroid hormone
regulation of circulating fibrinogen levels
adenylate cyclase activator activity
negative regulation of vesicle fusion
regulatory region RNA binding
signaling receptor activity
molecular transducer activity
axonogenesis
inorganic ion transmembrane transport
regulation of secretion
response to lipid
inorganic cation transmembrane transport
monoatomic cation transmembrane transport
intracellular chemical homeostasis
regulation of defense response
1-19 Red
pancreatic A cell development
rhombomere morphogenesis
cellular response to copper ion starvation
diamine oxidase activity
histamine oxidase activity
methylputrescine oxidase activity
propane-1,3-diamine oxidase activity
response to azide
cellular response to azide
response to copper ion starvation
transmembrane signaling receptor activity
signaling receptor activity
molecular transducer activity
G protein-coupled receptor signaling pathway
side of membrane
regulation of cell-cell adhesion
calcium ion binding
positive regulation of cell adhesion
inflammatory response
organic hydroxy compound metabolic process
3-1 Mosaic
cGMP-stimulated cyclic-nucleotide phosphodiesterase activity
spermidine transport
regulation of retina development in camera-type eye
spermidine transmembrane transporter activity
CNTFR-CLCF1 complex
parathyroid hormone receptor activity
regulation of gap junction assembly
positive regulation of skeletal muscle tissue regeneration
histone deacetylase regulator activity
positive regulation of gap junction assembly
regulation of secretion
regulation of protein transport
axonogenesis
regulation of establishment of protein localization
inorganic ion transmembrane transport
side of membrane
axon development
regulation of vesicle-mediated transport
synapse organization
regulation of anatomical structure size
2-10 Red
snoRNP binding
platelet dense tubular network membrane
adenine transport
thymine transport
DNA double-strand break attachment to nuclear envelope
deoxyribonuclease I activity
platelet dense tubular network
re-entry into mitotic cell cycle
CNTFR-CLCF1 complex
RNA 2'-O-methyltransferase activity
transmembrane signaling receptor activity
receptor complex
calcium ion binding
side of membrane
cell-substrate adhesion
calcium ion homeostasis
postsynaptic membrane
basal plasma membrane
regulation of T cell activation
basal part of cell
3-2 Mosaic
post-embryonic body morphogenesis
trigeminal ganglion development
T cell antigen processing and presentation
negative regulation of ceramide biosynthetic process
Kv4.2-KChIP2 channel complex
benzaldehyde dehydrogenase (NAD+) activity
benzaldehyde dehydrogenase [NAD(P)+] activity
3-chloroallyl aldehyde dehydrogenase activity
UDP-N-acetylgalactosamine metabolic process
troponin complex
regulation of secretion
inorganic cation transmembrane transport
inorganic ion transmembrane transport
monoatomic cation transmembrane transport
modulation of chemical synaptic transmission
axon development
regulation of neuron projection development
regulation of vesicle-mediated transport
regulation of MAPK cascade
regulation of anatomical structure size

Figure S4

### Enrichment analysis in TRRUST

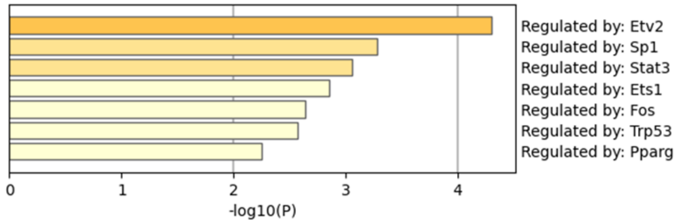


Figure S5

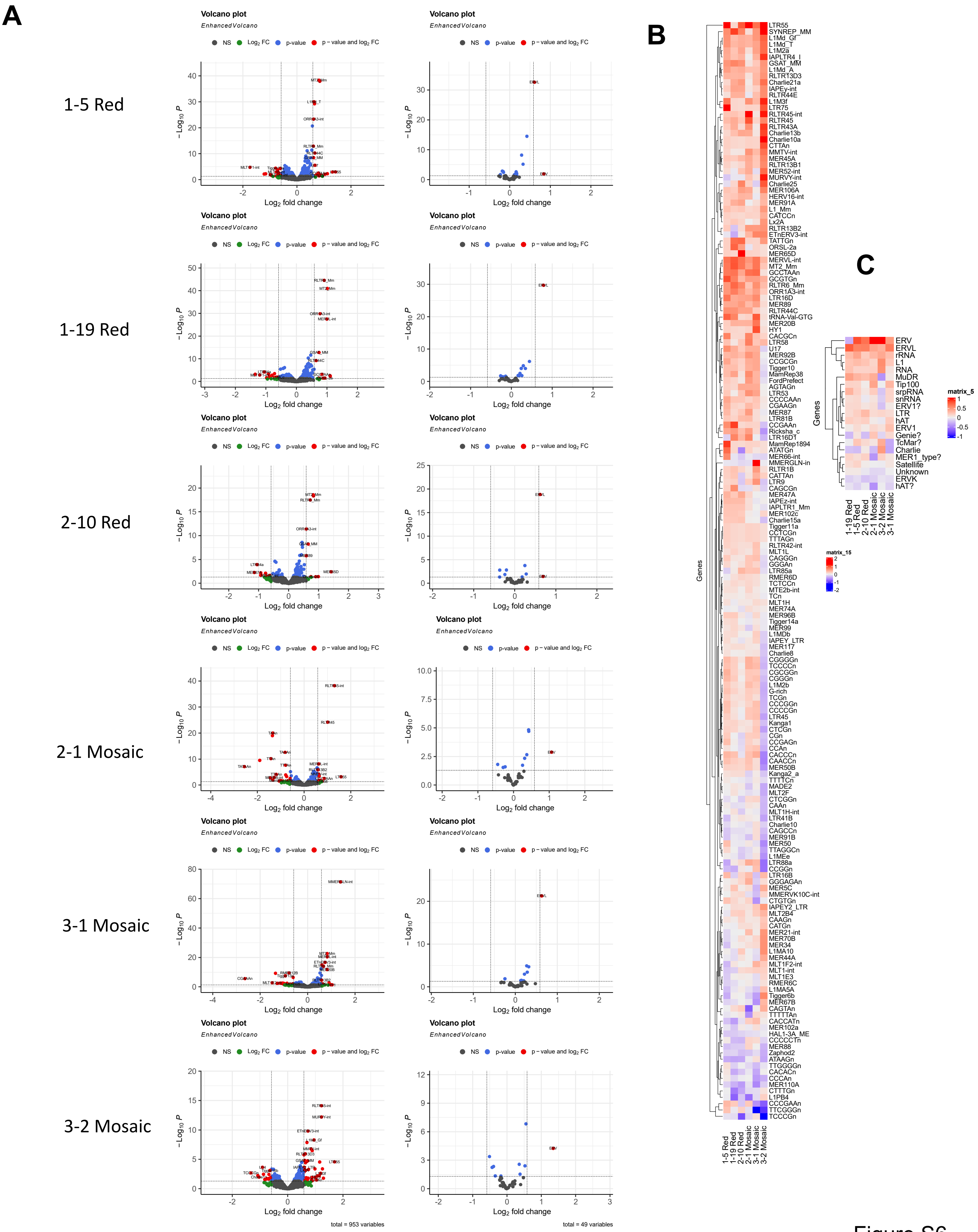

A
B
C
1-5 Red
1-19 Red
2-10 Red
2-1 Mosaic
3-1 Mosaic
3-2 Mosaic
Volcano plot
EnhancedVolcano
NS
Log2 FC
p-value
p – value and log2 FC
−Log10 P
Log2 fold change
total = 953 variables
total = 49 variables
Genes
matrix_15
matrix_5
LTR55
SYNREP_MM
L1Md_Gf
L1Md_T
L1M2a
IAPLTR4_I
GSAT_MM
L1Md_A
RLTR13D3
Charlie21a
IAPEy-int
RLTR44E
L1M3f
LTR75
RLTR45-int
RLTR45
RLTR43A
Charlie13b
Charlie10a
CTTAn
MMTV-int
MER45A
RLTR13B1
MER52-int
MURVY-int
Charlie25
MER106A
HERV16-int
MER91A
L1_Mm
CATCCn
Lx2A
RLTR13B2
ETnERV3-int
TATTGn
ORSL-2a
MER65D
MERVL-int
MT2_Mm
GCCTAAn
GCGTGn
RLTR6_Mm
ORR1A3-int
LTR16D
MER89
RLTR44C
tRNA-Val-GTG
MER20B
HY1
CACGCn
LTR58
U17
MER92B
CCGCGn
Tigger10
MamRep38
FordPrefect
AGTAGn
LTR53
CCCCAAn
CGAAGn
MER87
LTR81B
CCGAAn
Ricksha_c
LTR16D1
MamRep1894
ATATGn
MER66-int
MMERGLN-int
RLTR1B
CATTAn
LTR9
CAGCGn
MER47A
IAPEz-int
IAPLTR1_Mm
MER102c
Charlie15a
Tigger11a
CCTCGn
TTTAGn
RLTR42-int
MLT1L
CAGGGn
GGGAn
LTR85a
RMER6D
TCTCCn
MTE2b-int
TCn
MLT1H
MER74A
MER96B
Tigger14a
MER99
L1MDb
IAPEY_LTR
MER117
Charlie8
CGGGGn
TCCCCn
CGCGGn
CGGGn
L1M2b
G-rich
TCGn
CCCGGn
CCCCGn
LTR45
Kanga1
CTCGn
CGn
CCGAGn
CCAn
CACCCn
CAACCn
MER50B
Kanga2_a
TTTTCn
MADE2
MLT2F
CTCGGn
CAAn
MLT1H-int
LTR41B
Charlie10
CAGCCn
MER91B
MER50
TTAGGCn
L1MEe
LTR88a
CCGGn
LTR16B
GGGAGAn
MER5C
MMERVK10C-int
CTGTGn
IAPEY2_LTR
MLT2B4
CAAGn
CATGn
MER21-int
MER70B
MER34
L1MA10
MER44A
MLT1F2-int
MLT1-int
MLT1E3
RMER6C
L1MA5A
Tigger6b
MER67B
CAGTAn
TTTTTAn
CACCATn
MER102a
HAL1-3A_ME
CCCCCTn
MER88
Zaphod2
ATAAGn
TTGGGGn
CACACn
CCCAn
MER110A
CTTTGn
L1PB4
CCCGAAn
TTCGGGn
TCCCGn
ERV
ERVL
rRNA
L1
RNA
MuDR
Tip100
srpRNA
snRNA
ERV1?
LTR
hAT
ERV1
Genie?
TcMar?
Charlie
MER1_type?
Satellite
Unknown
ERVK
hAT?
1-19 Red
1-5 Red
2-10 Red
2-1 Mosaic
3-2 Mosaic
3-1 Mosaic
1-5 Red
1-19 Red
2-10 Red
2-1 Mosaic
3-1 Mosaic
3-2 Mosaic


Figure S6

Day 12 EB Differentiation and Segmentation

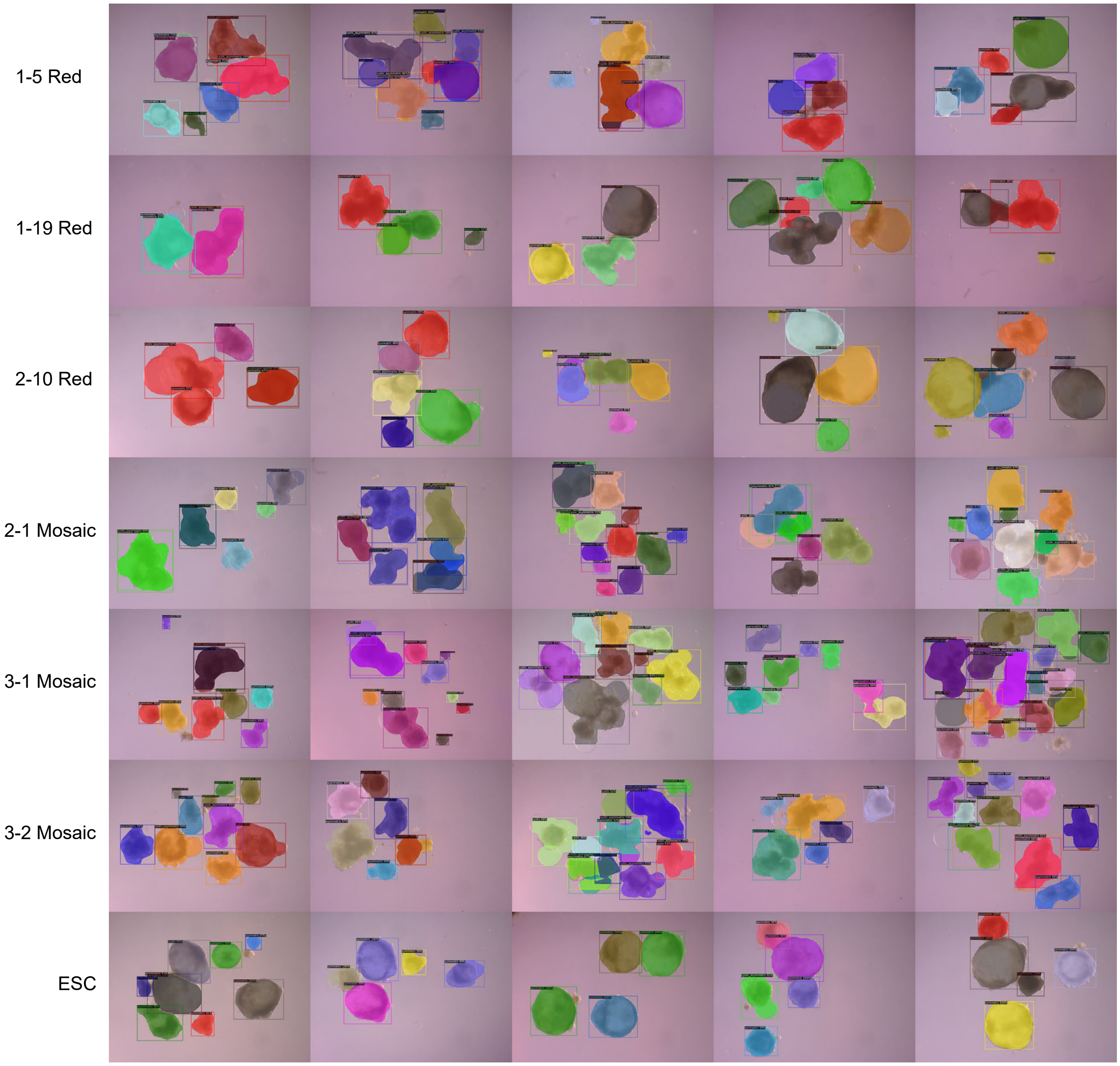


Figure S7A

Day 12 EB Differentiation

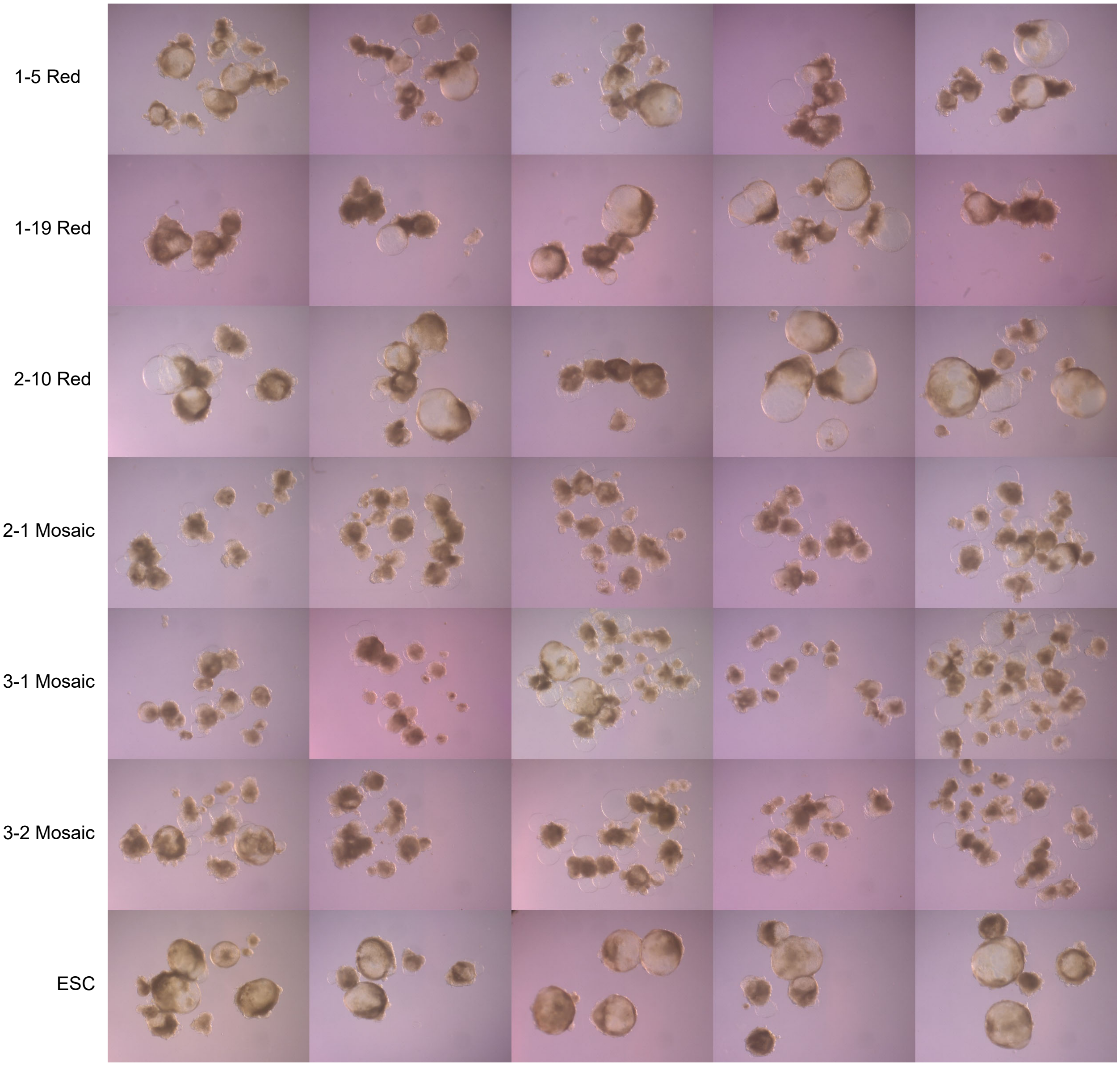


Figure S7B

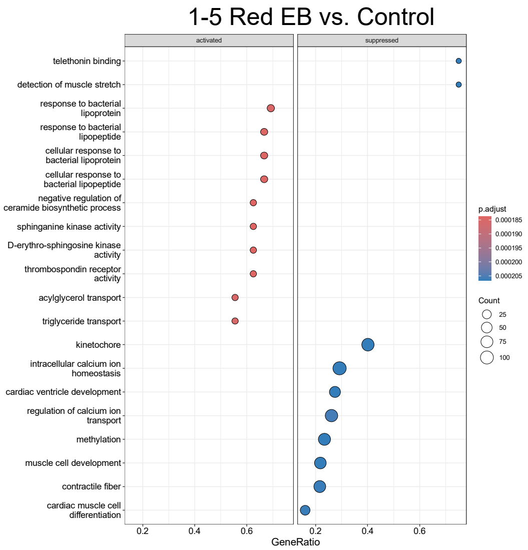

1-5 Red EB vs. Control
activated
suppressed
telethonin binding
detection of muscle stretch
response to bacterial lipoprotein
response to bacterial lipopeptide
cellular response to bacterial lipoprotein
cellular response to bacterial lipopeptide
negative regulation of ceramide biosynthetic process
sphinganine kinase activity
D-erythro-sphingosine kinase activity
thrombospondin receptor activity
acylglycerol transport
triglyceride transport
kinetochore
intracellular calcium ion homeostasis
cardiac ventricle development
regulation of calcium ion transport
methylation
muscle cell development
contractile fiber
cardiac muscle cell differentiation
0.2
0.4
0.6
0.2
0.4
0.6
GeneRatio
p.adjust
0.000185
0.000190
0.000195
0.000200
0.000205
Count
25
50
75
100


Figure S8

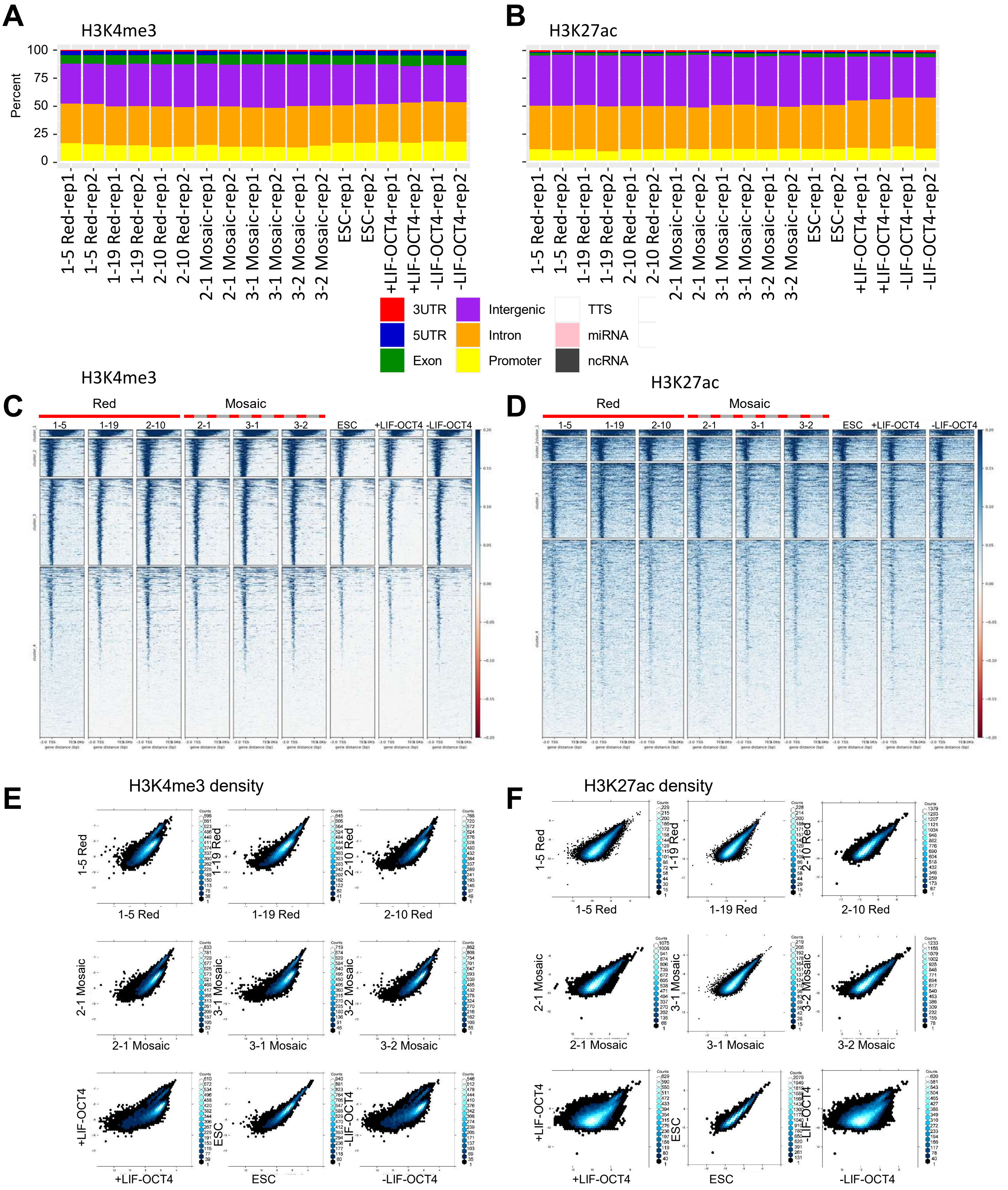
A
H3K4me3
Percent
100
75
50
25
0
1-5 Red-rep1
1-5 Red-rep2
1-19 Red-rep1
1-19 Red-rep2
2-10 Red-rep1
2-10 Red-rep2
2-1 Mosaic-rep1
2-1 Mosaic-rep2
3-1 Mosaic-rep1
3-1 Mosaic-rep2
3-2 Mosaic-rep1
3-2 Mosaic-rep2
ESC-rep1
ESC-rep2
+LIF-OCT4-rep1
+LIF-OCT4-rep2
-LIF-OCT4-rep1
-LIF-OCT4-rep2
B
H3K27ac
3UTR
5UTR
Exon
Intergenic
Intron
Promoter
TTS
miRNA
ncRNA
C
H3K4me3
Red
Mosaic
1-5
1-19
2-10
2-1
3-1
3-2
ESC
+LIF-OCT4
-LIF-OCT4
D
H3K27ac
E
H3K4me3 density
1-5 Red
1-19 Red
2-10 Red
2-1 Mosaic
3-1 Mosaic
3-2 Mosaic
+LIF-OCT4
ESC
-LIF-OCT4
F
H3K27ac density

Figure S9

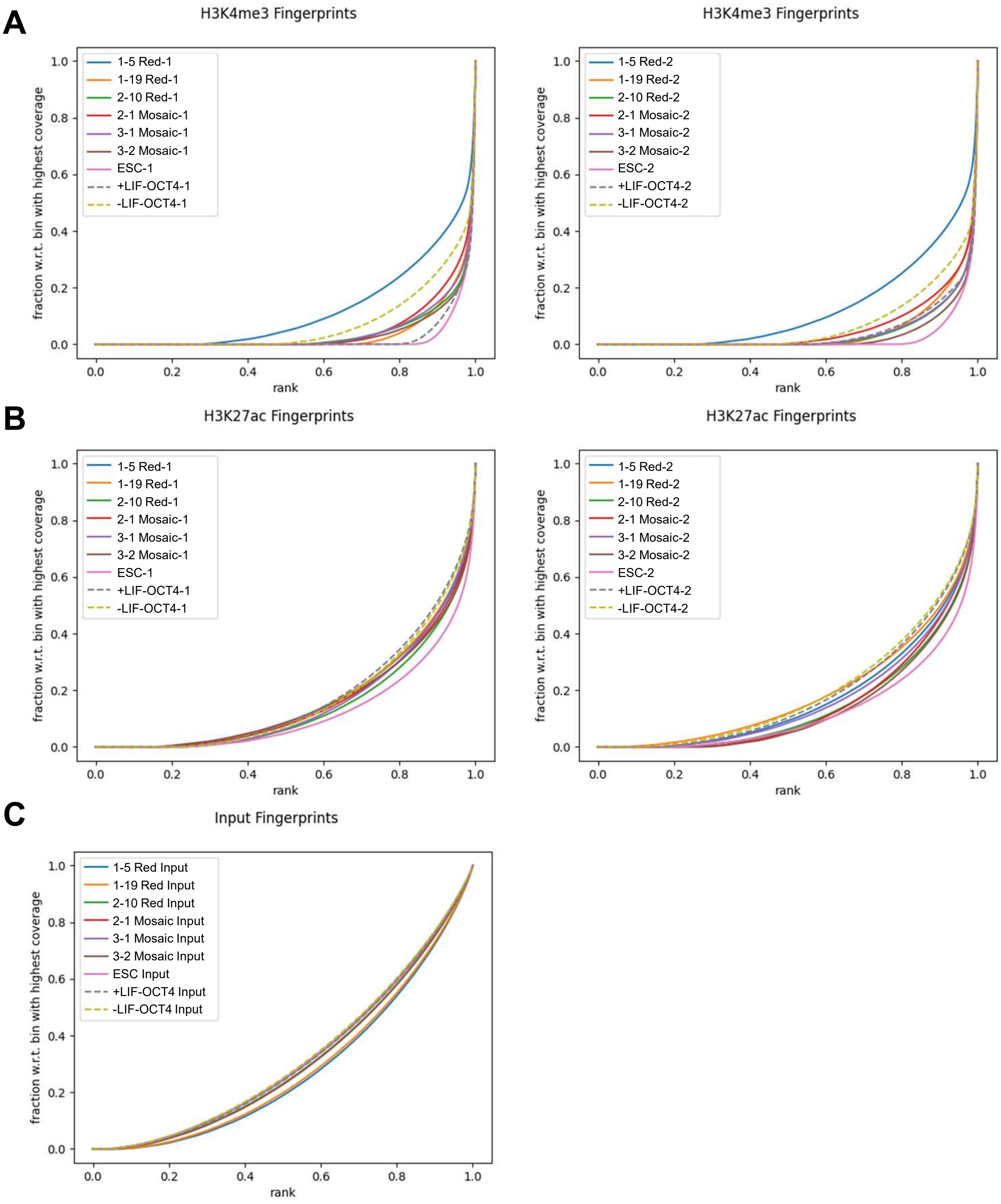
A
H3K4me3 Fingerprints
1-5 Red-1
1-19 Red-1
2-10 Red-1
2-1 Mosaic-1
3-1 Mosaic-1
3-2 Mosaic-1
ESC-1
+LIF-OCT4-1
-LIF-OCT4-1
fraction w.r.t. bin with highest coverage
rank
H3K4me3 Fingerprints
1-5 Red-2
1-19 Red-2
2-10 Red-2
2-1 Mosaic-2
3-1 Mosaic-2
3-2 Mosaic-2
ESC-2
+LIF-OCT4-2
-LIF-OCT4-2
fraction w.r.t. bin with highest coverage
rank
B
H3K27ac Fingerprints
1-5 Red-1
1-19 Red-1
2-10 Red-1
2-1 Mosaic-1
3-1 Mosaic-1
3-2 Mosaic-1
ESC-1
+LIF-OCT4-1
-LIF-OCT4-1
fraction w.r.t. bin with highest coverage
rank
H3K27ac Fingerprints
1-5 Red-2
1-19 Red-2
2-10 Red-2
2-1 Mosaic-2
3-1 Mosaic-2
3-2 Mosaic-2
ESC-2
+LIF-OCT4-2
-LIF-OCT4-2
fraction w.r.t. bin with highest coverage
rank
C
Input Fingerprints
1-5 Red Input
1-19 Red Input
2-10 Red Input
2-1 Mosaic Input
3-1 Mosaic Input
3-2 Mosaic Input
ESC Input
+LIF-OCT4 Input
-LIF-OCT4 Input
fraction w.r.t. bin with highest coverage
rank

Figure S10

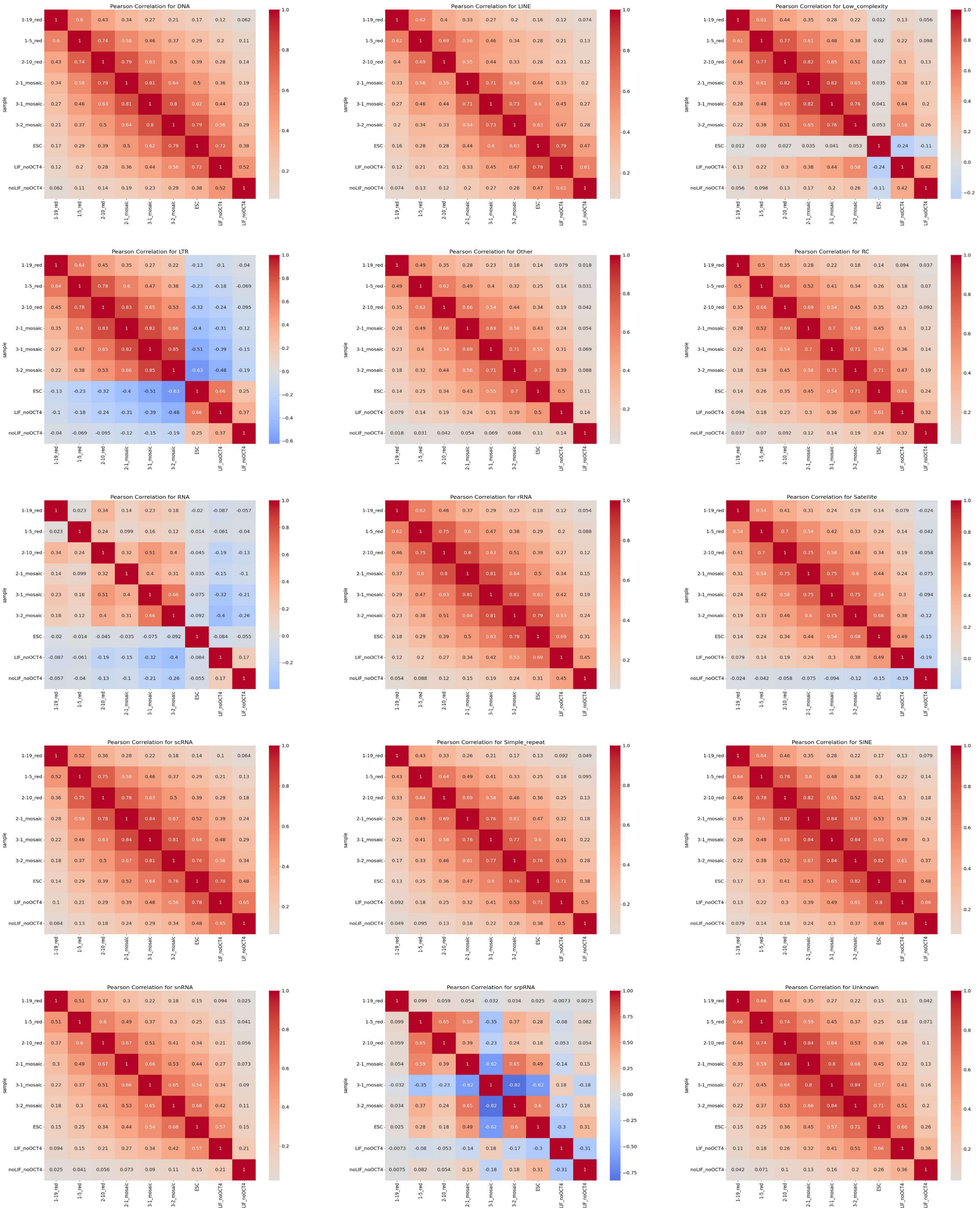
Pearson Correlation for DNA
Pearson Correlation for LINE
Pearson Correlation for Low_complexity
Pearson Correlation for LTR
Pearson Correlation for Other
Pearson Correlation for RC
Pearson Correlation for RNA
Pearson Correlation for rRNA
Pearson Correlation for Satellite
Pearson Correlation for scRNA
Pearson Correlation for Simple_repeat
Pearson Correlation for SINE
Pearson Correlation for snRNA
Pearson Correlation for srpRNA
Pearson Correlation for Unknown
sample
1-19_red
1-5_red
2-10_red
2-1_mosaic
3-1_mosaic
3-2_mosaic
ESC
LIF_noOCT4
noLIF_noOCT4

Figure S11

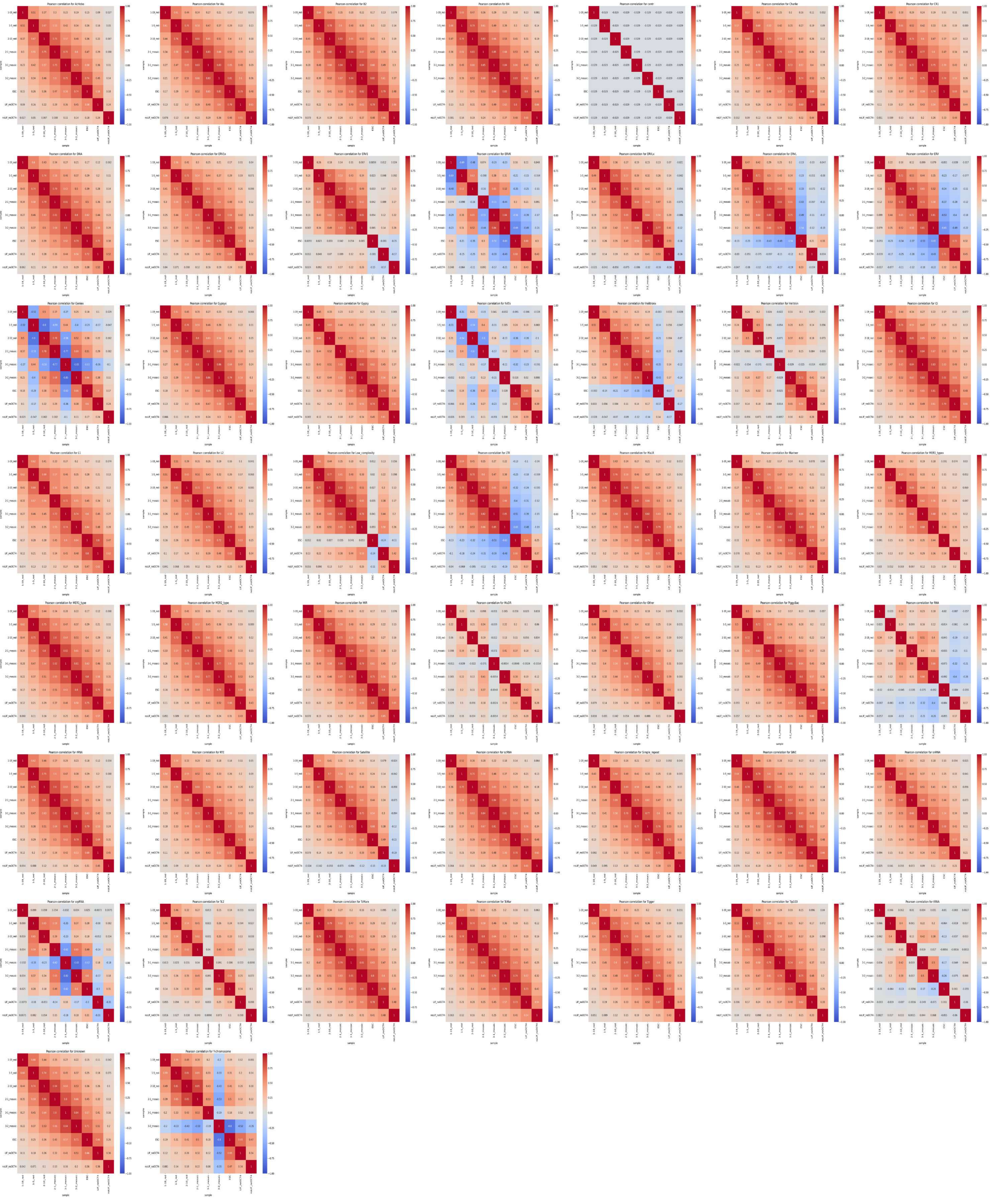

Figure S12
